# Inclusion of JNK-independent drugs within multi-agent chemotherapy improves response in relapsed high-risk neuroblastoma

**DOI:** 10.1101/2025.04.26.650760

**Authors:** Jeremy Z.R. Han, Monica Phimmachanh, Jordan F. Hastings, King Ho Leong, Boaz H. Ng, Jenny Ni, Angela Fontaine-Titley, Antonia L. Cadell, Yolande EI O’Donnell, Misaki Clearwater, Alvin Kamili, Michelle Haber, Murray Norris, Paul Timpson, Toby N. Trahair, Jamie I. Fletcher, Dirk Fey, Walter Kolch, Sharissa L. Latham, David R. Croucher

## Abstract

The acquisition of a chemoresistant state underlies poor prognosis in many cancers, including neuroblastoma. We previously demonstrated that heterogeneity in apoptosis induction through c-Jun N-Terminal Kinase (JNK) promotes a form of non-genetic chemoresistance in neuroblastoma observable at both patient and single-cell levels. As the maintenance of this JNK-impaired state in the relapse setting is a significant barrier to the efficacy of many standard-of-care chemotherapy drugs, we combined a mechanistic, mathematical model of JNK activation with a paediatric focused drug screen and identified approved oncology drugs capable of inducing apoptosis in a JNK-independent manner. Functional genomics further revealed that synergy between these JNK-independent drugs and standard-of-care chemotherapies emerged from differential utilisation of apoptotic network components, rather than from their direct mechanistic targets. Efficacy studies with PDX models also confirmed that including a JNK-independent drug within existing chemotherapy backbones significantly improved response in the relapse setting, where new approaches are urgently needed.

**One Sentence Summary:** Drug combinations utilising differential sets of network components can overcome the apoptotic impaired state of relapsed high-risk neuroblastoma.

## Introduction

Neuroblastoma is the most common solid paediatric cancer outside of the brain, accounting for 7-8% of childhood cancers (1). These tumours are derived from neural crest progenitor cells and manifest along the sympathetic nervous system, most commonly in the adrenal medulla and the sympathetic ganglia (2). In low and intermediate-risk neuroblastoma, surgical resection of the primary tumour is usually sufficient to achieve complete remission, with survival rates approaching 90-98% (2, 3). Unfortunately, most patients are diagnosed with high-risk neuroblastoma (HRNB), an aggressive and metastatic disease. Despite multi-modal treatment including intense multi-agent chemotherapy, surgery, radiotherapy and immunotherapy, less than 50% of HRNB patients are cured. In fact, 15% of HRNB patients do not respond to chemotherapy, while a further half will relapse following an initial response (4, 5). For these patients with chemoresistant disease, the 5-year overall survival is a dire 10% (6, 7).

Another factor in the poor overall prognosis of HRNB is the lack of targeted therapies. HRNB is highly heterogeneous, and outside of *MYCN* amplification and *ALK* activating mutations, disease is driven by a combination of multiple low frequency mutations and deleterious chromosomal events (8, 9). Consequently, multi-agent chemotherapy still forms the backbone of existing standard-of-care for patients with HRNB. While regimens vary around the world, neoadjuvant (induction) therapy typically consists of multiple cycles of a number of chemotherapy drugs, including various combinations of topotecan, cyclophosphamide, cisplatin, carboplatin, etoposide, vincristine and doxorubicin (10–12). Following this initial treatment, patients with refractory, or relapsed disease, will undergo salvage therapy with different combination treatments including TVD (topotecan, vincristine and doxorubicin) (13), IT/TEMIRI (irinotecan and temozolomide) (14) and TOTEM (topotecan and temozolomide) (15).

These combinations of mechanistically different chemotherapy drugs have been a cornerstone of cancer care for many decades, often considered an effective way of combatting the chemoresistance that arises from patient heterogeneity, intra-tumour heterogeneity and the adaptive dysregulation of signalling pathways (16, 17). However, the mechanisms through which these drugs may potentially combine to produce a beneficial clinical response is not always considered, and often not well understood, particularly within the highly drug-resistant state of relapsed or refractory disease.

Many studies have focused on the potential genetic mechanisms underlying the resistant state of HRNB, only occasionally identifying therapeutically actionable mutations emerging within individual relapsed tumours (8, 9). Therefore, in order to develop more broadly applicable therapeutic options, we previously focused on the mechanisms underpinning non-genetic chemoresistance in neuroblastoma by using patient-specific mathematical modelling of drug-induced apoptotic signalling (18). This approach demonstrated that an impaired ability to activate apoptosis through c-Jun N-Terminal Kinase (JNK) - a pathway required the induction of apoptosis by many standard-of-care chemotherapy drugs - was significantly associated with poor overall survival for HRNB patients (18). By further adapting this model to simulate single-cell drug response dynamics, we also identified a stochastic population of JNK-impaired cells undergoing positive selection, and stabilization, within relapsed tumours (19). While these innately chemoresistant cells could be re-sensitized by the pre-treatment of naïve tumours with the Histone Deacetylase (HDAC) inhibitor vorinostat, relapsed tumours were resistant to this priming regimen.

Alternative approaches to overcoming this JNK-impaired state within relapsed HRNB are required to extend this paradigm. Therefore, here we combined a stratified panel of neuroblastoma cell lines with a pediatric-focused drug screen in order to identify approved oncology drugs capable of initiating apoptosis in a JNK-independent manner. Further characterization of the mechanism of action for these drugs, along with an apoptosis-focused functional genomics approach, identified fundamental requirements for the selection of synergistic drug pairs in HRNB. This analysis demonstrated that the emergence of synergy was based upon the differential utilization of apoptotic network components by each drug, rather than the combination of drugs targeting different biochemical pathways. Accordingly, proof-of-concept experiments with both *ex vivo* and *in vivo* patient-derived xenograft (PDX) models demonstrate the utility of introducing JNK-independent drugs into salvage treatment schedules. These findings have important implications for the existing standard-of-care chemotherapy combinations used to treat relapsed HRNB, suggesting that incorporating JNK-independent drugs could improve response rates and outcomes for patients with relapsed HRNB.

## Results

### Identification of JNK-Independent Drugs

As a key mediator of stress-induced signalling, the JNK pathway is required for chemotherapy induced apoptosis within a number of tumour types (20). We have previously demonstrated that JNK activation is required in response to standard-of-care chemotherapies in neuroblastoma, extending this observation to the development of a dynamic model of JNK signalling used to produce both prognostically significant patient-specific predictions of drug response (18) and to simulate single-cell apoptotic JNK-signalling (19). Here, we leveraged this model to identify drugs capable of activating apoptosis without the need for JNK activation. This was achieved through the stratification of a panel of neuroblastoma cell lines with varying abilities to activate JNK in response to treatment with JNK-dependent chemotherapies such as vincristine (**Fig. 1A**). This modelling analysis was performed using the relative expression levels of the JNK network components; including JNK1/2, mitogen-activated protein kinase (MAPK) kinase 4 (MKK4), MAPK kinase 7 (MKK7), MAPK kinase kinase 20 (ZAK), and phospho-Akt (Ser473). As we have performed previously (18, 19), these relative expression levels were used as parameters within our ordinary differential equation (ODE) model of JNK activation to produce predictions of JNK activation for each cell line (**Fig. 1B**). Importantly, these simulations again demonstrated that predicted JNK activity correlated strongly with the observed JNK phosphorylation levels measured by Western blotting following vincristine treatment (100 nM, 2 h. R^2^ = 0.9239) (**Fig. 1C**). In addition, high content imaging-based readouts of Caspase-3/7 activity were also measured for each cell line in response to vincristine (100 nM, 24 h) (**Fig 1D**). This direct readout of apoptosis also correlated strongly with the predicted ability of each cell line to activate JNK (R^2^ = 0.9049), demonstrating that our predictive model provided a high-confidence stratification for this JNK-dependent chemotherapy drug.

**Figure 1:**
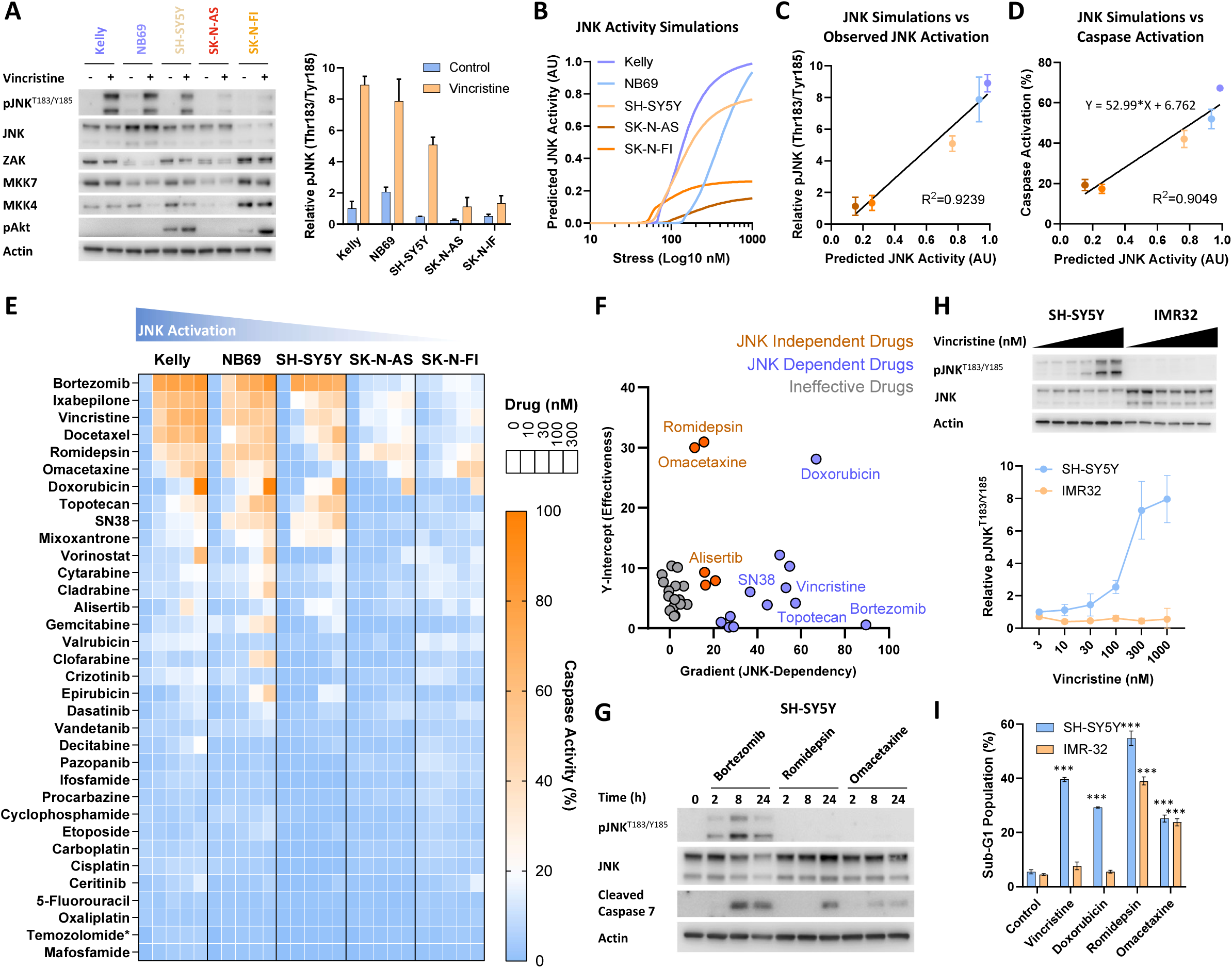
Identification of JNK-independent drugs. (**A**) Western blot analysis of neuroblastoma cell lines treated with vincristine (300 nM, 2 h), using the antibodies indicated (Mean ± SD, n=3). (**B**) Quantified values from A were used to perform simulations of predicted JNK activation in each cell line. (**C**) Correlation between the observations of vincristine-induced JNK activity in A and the predicted simulations in B. (**D**) Correlation between the predicted JNK activity simulations in B and the percentage of cells with caspase activity after staining with NucView 488 (MFI > 20), measured by high-content imaging following vincristine treatment (300 nM, 24 h). (**E**) The percentage of cells with caspase activity (MFI > 20) following treatment with the drugs and doses indicated (24 h). * indicates an alternative dosage range of 0, 100, 300, 1000, 3000 nM. (**F**) Correlation between cell line simulations of JNK-activity in panel B and the caspase activity measured for all drugs in E. For each drug, line fitting was performed as in panel D, the gradient and extrapolated y-intercept calculated and correlated within an XY plot. (**G**) Western blot analysis with the antibodies indicated in the SH-SY5Y cell line following treatment with the drugs indicated (300 nM). (**H**) Western blot analysis with the antibodies indicated in the SH-SY5Y and IMR32 cell lines following treatment with increasing doses of vincristine (3, 10, 30, 100, 300, 1000 nM. 2 h) (Mean ± SD, n=3). (**I**) Analysis of the sub-G1 population measured by flow cytometry in the SH-SY5Y and IMR32 cell lines following treatment with the drugs indicated (300 nM, 24 h) and propidium iodide staining (Mean ± SD, n=3). *** p<0.001.

Exploiting this, we sought to find chemotherapies for which our simulations of JNK activation did not correlate with apoptotic output, suggesting a potential lack of JNK dependency for these specific drugs. To achieve this, these stratified cell lines were treated with an expanded panel of FDA-approved drugs, which all had available paediatric safety data (**Table S1**). High content imaging with a Caspase 3/7 fluorescent substrate was also performed, to quantify the percentage of cells undergoing apoptosis in response to increasing doses of each drug (**Fig. 1E**). For each drug, caspase activation in each cell line was correlated to the respective JNK activity simulations, as already performed for vincristine (**Fig. 1D**). From the resultant plots, linear fitting was performed and values were extracted from the standard slope intercept form (*y = mx + b*). For this analysis, the gradient (*m*) of the fitted line was assumed to reflect the relative JNK-dependency of each drug, with JNK-dependent drugs expected to exhibit a steeper gradient and JNK-independent drugs expected to be flatter. In addition, the extrapolated Y-intercept value (*b*) was used as a readout of the broad effectiveness of each drug across the cell line panel. By correlating these two outputs for the whole panel, this dataset clearly differentiated between drugs that appeared to be highly JNK-dependent (Steep gradient, low Y-intercept) and those that were potentially JNK-independent (Flat gradient, higher Y-intercept) (**Fig. 1F**). In line with expectations from our previous studies (18, 19), many of the current standard-of-care drugs for HRNB were identified as JNK dependent, including the topoisomerase I inhibitors topotecan and SN38 (active metabolite of irinotecan), the topoisomerase II inhibitor doxorubicin (21) and the microtubule destabiliser vincristine. On top of these established HRNB drugs, bortezomib, a proteasome inhibitor used in T-cell acute lymphoblastic leukemia and lymphoma (22), was also identified as potentially the most JNK-dependent drug in this panel.

Importantly, this analysis also identified two notable outliers, romidepsin and omacetaxine, which were both broadly effective and exhibited a poor correlation with the JNK activity simulations. Omacetaxine is a protein translation inhibitor currently used to treat chronic myeloid leukaemia, while romidepsin is an HDAC inhibitor currently used to treat T-cell lymphoma (23). Romidepsin is also known to induce caspase-dependent apoptosis in neuroblastoma cells (24, 25) and has already completed successful Phase I clinical trials in paediatric solid tumours (26). To confirm the potential insights from this mechanistic drug screen, we further investigated the ability of bortezomib (JNK-dependent), along with romidepsin and omacetaxine (JNK-independent), to activate both JNK signalling and apoptosis by Western blotting (**Fig. 1G**). As expected, bortezomib strongly induced both JNK phosphorylation and caspase cleavage, while romidepsin and omacetaxine were capable of inducing caspase cleavage but did so without any detectable JNK phosphorylation.

Further validation of these findings was also provided using the IMR32 cell line, which does not activate JNK in response to vincristine due to alterations in expression of multiple components of the JNK network (18). A direct comparison of vincristine-induced JNK phosphorylation in both the SH-SY5Y and IMR32 cell lines confirmed the inability of the IMR32 line to activate JNK (**Fig 1H**). We therefore investigated the induction of apoptosis in both cell lines using an orthogonal readout - quantification of the sub-G1 cell population following propidium iodide staining as an indication of DNA fragmentation and late-stage apoptosis (**Fig. 1I**). While the SH-SY5Y cell line was capable of undergoing a significant induction of apoptosis in response to both JNK-dependent (vincristine and doxorubicin) and JNK-independent (romidepsin and omacetaxine) drugs, the JNK-impaired IMR32 cell line only responded to the JNK-independent drugs.

Together, these observations demonstrated that omacetaxine and romidepsin do not require JNK activity to activate apoptosis in neuroblastoma cells, prompting further investigation into their specific mechanisms-of-action and their potential to combine with existing JNK-dependent standard-of-care chemotherapy drugs to improve response in the setting of JNK-impaired relapse HRNB.

### Correlating target engagement with apoptosis induction

To further investigate the link between mechanism-of-action and the induction of apoptosis in neuroblastoma cells, we sought to directly correlate cellular target engagement with caspase cleavage for each of these drugs. For these experiments, our underlying hypothesis suggested that there would be a weak correlation between target engagement and apoptosis induction for JNK-dependent drugs, with the downstream bottleneck of JNK activation acting as the major factor controlling cell death, rather than a limitation within the direct, initial action of the drugs upon the cell.

Both of the JNK-dependent, standard-of-care drugs topotecan and doxorubicin interfere with topoisomerase I/II function, respectively, and result in the accumulation of DNA damage and the activation of apoptosis (27). Therefore, immunofluorescence with a γH2A.X antibody was used as a readout of DNA damage after treatment with either doxorubicin or topotecan (100 nM; 24 hours), which results in punctate nuclear foci following both treatments (**Fig. 2A,C**). To correlate this readout of drug activity with the previous caspase activation data (**Fig. 1E**), a dose response was performed with each cell line and the relative fluorescence intensity of the γH2A.X foci quantified. While both drug treatments significantly increased the intensity of γH2A.X foci across all lines (**Fig. S1A,B**), there was a poor correlation between the formation of these foci and caspase activation (R^2^ = 0.3454 for doxorubicin, R^2^ = 0.1147 for topotecan (**Fig. 2B,D**).

**Figure 2:**
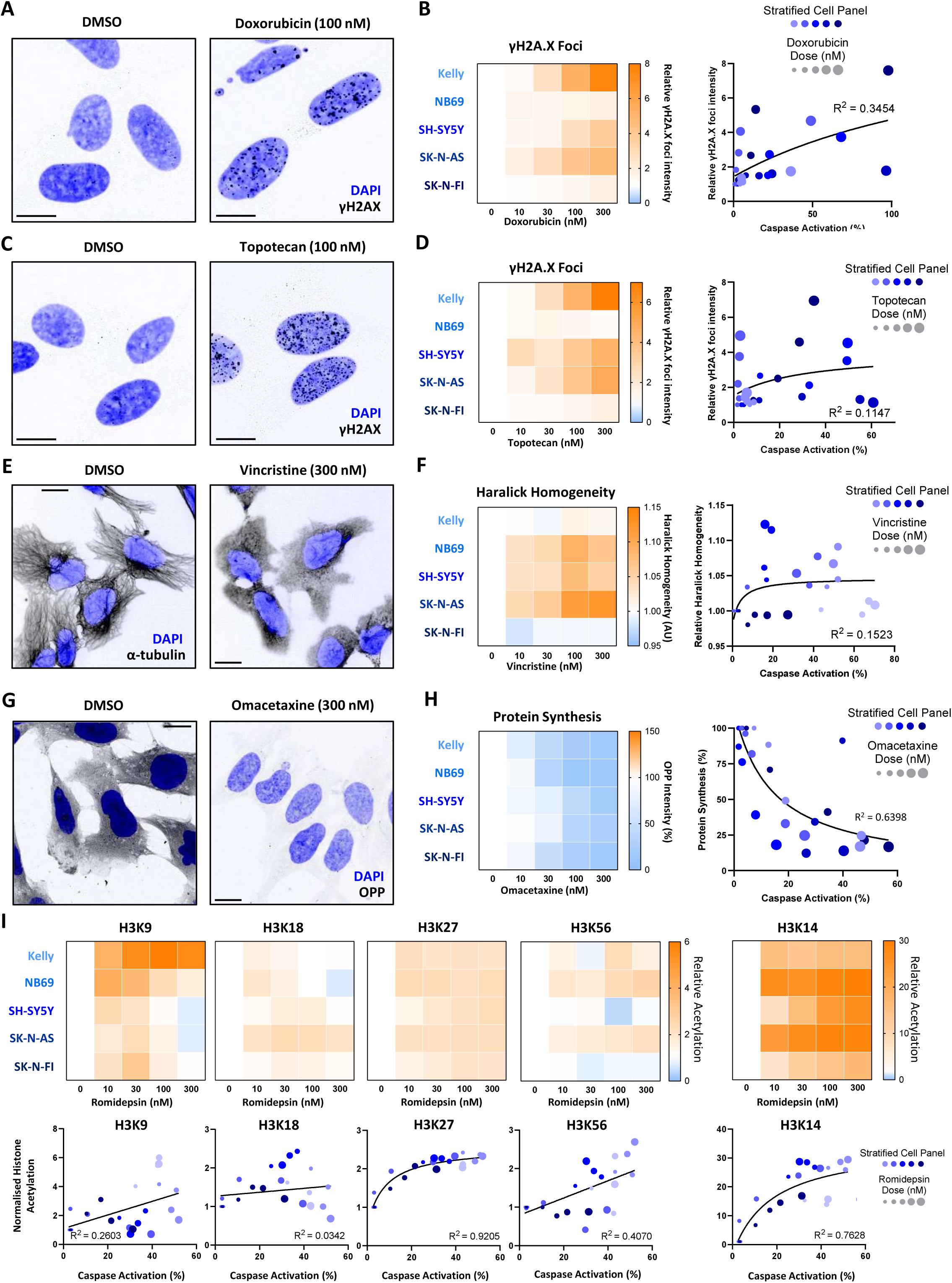
Correlating target engagement with apoptosis induction. (**A**) Confocal fluorescence imaging of γH2A.X antibody and DAPI staining in the SH-SY5Y cell line following treatment with doxorubicin (100 nM. 24 h) or a DMSO control. (**B**) Quantification of γH2A.X foci intensity by high-content imaging in the cell lines indicated following doxorubicin treatment at the dosages indicated (24 h. Mean, n>1000) and the correlation with doxorubicin induced caspase activity in Figure 1E. (**C**) Confocal fluorescence imaging of γH2A.X antibody and DAPI staining in the SH-SY5Y cell line following treatment with topotecan (100 nM. 24 h) or a DMSO control. (**D**) Quantification of γH2A.X foci intensity by high content imaging in the cell lines indicated following topotecan treatment at the dosages indicated (24 h. Mean, n>1000) and the correlation with topotecan induced caspase activity in Figure 1E. (**E**) Confocal fluorescence imaging of α-tubulin antibody and DAPI staining in the SK-N-AS cell line following treatment with vincristine (300 nM. 4 h) or a DMSO control. (**F**) Quantification of Haralick homogeneity by high content imaging in the cell lines indicated following vincristine treatment at the dosages indicated (4 h. Mean, n>1000) and the correlation with vincristine induced caspase activity in Figure 1E. (**G**) Confocal fluorescence imaging of OPP and DAPI staining in the SH-SY5Y cell line following treatment with omacetaxine (300 nM. 24 h) or a DMSO control. (**H**) Quantification of OPP intensity by high content imaging in the cell lines indicated following omacetaxine treatment at the dosages indicated (24 h. Mean, n>1000) and the correlation with vincristine induced caspase activity in Figure 1E. (**I**) Quantification of relative histone acetylation at the sites indicated by high content imaging of antibody staining in the cell lines indicated following romidepsin treatment at the dosages indicated (16 h. Mean, n>1000) and the correlation with romidepsin induced caspase activity in Figure 1E. All scale bars are 10 µm.

The JNK-dependent chemotherapy drug vincristine acts by destabilising microtubules, which are integral to cell motility, mitosis and cell shape. Therefore, as expected, vincristine treatment (300 nM; 24 hours) resulted in a clear reduction in distinct, linear α-tubulin fibres within the cytosol (**Fig. 2E**). Quantification of this α-tubulin directionality was performed across the cell line panel using the Haralick texture method (28). While there was a significant dose-dependent increase in Haralick homogeneity for all lines, apart from the Kelly and SK-N-FI lines (**Fig. S1C**), this also correlated poorly with caspase activation (R^2^ = 0.1523) (**Fig. 2F**).

The JNK-independent drug omacetaxine is known to directly interact with the ribosomal-A site, preventing initial elongation during protein synthesis, thereby inhibiting protein translation (29). Therefore, to quantify drug action, a fluorescent analogue of puromycin, O-propargyl-puromycin (OPP), was utilised as a readout of protein synthesis. Following omacetaxine treatment (300 nM; 24 hours) there was a marked reduction in OPP and thus protein synthesis (**Fig. 2G**). Quantifying the dose response of omacetaxine treatment also resulted in a significant decrease in protein synthesis across all lines (**Fig S1D**), although in contrast to the JNK-dependent drugs, there was a good correlation between the inhibition of protein synthesis and caspase activation (R^2^ = 0.6398) (**Fig. 2H**).

The second JNK-independent drug, romidepsin, is a Class I HDAC inhibitor and thereby impairs deacetylation at multiple histone residues (24, 30). Expectedly, treatment with romidepsin resulted in increased histone acetylation at all sites measured (**Fig. S1E,F**). In particular, romidepsin induced robust and significant increases acetylation at H3K27 and H3K14 acetylation across the entire cell panel, with a strong correlation between both of these acetylation sites and caspase activation (R^2^ = 0.9205 and R^2^ = 0.7628 respectively).

Taken together, this target engagement data supports the JNK-dependent and -independent nature of these drugs. The poor correlation between the induction of apoptosis and the direct action of doxorubicin, topotecan and vincristine reiterates the convergence of apoptotic signalling upon the JNK bottleneck for these drugs. Conversely, the strong correlation between target engagement and the induction of apoptosis for omacetaxine and romidepsin highlights their independence from JNK signalling and underpins their relatively increased cytotoxic activity within JNK-impaired cell lines.

### JNK-independent mechanisms of apoptotic signalling

As we have identified the ability of romidepsin and omacetaxine to induce apoptosis without the requirement of JNK signalling, we sought to further delineate their alternative mechanisms-of-action. Therefore, using a multiplexed bead-based proteomics platform we examined MAPK, PI3K/mTOR, DNA damage response and apoptosis regulation pathways within the NB69, SK-N-AS and SK-N-FI cell lines, following a time course of drug treatment with vincristine, romidepsin and omacetaxine (**Fig. 3A**), allowing an investigation of the mechanism underlying the action of each individual drug (**Fig. B**). This analysis again highlighted the JNK-impaired nature of the SK-N-AS and SK-N-FI cell lines in comparison to the NB69 line (**Fig. 3C**), along with the lack of JNK activation following treatment with romidepsin and omacetaxine. From a mechanistic viewpoint, we have previously demonstrated that vincristine induced JNK activation promotes the degradation of MCL-1 (19), which was also apparent within this dataset, occurring in proportion to the observed levels of JNK activation within each cell line (**Fig. 3B,D**).

**Figure 3:**
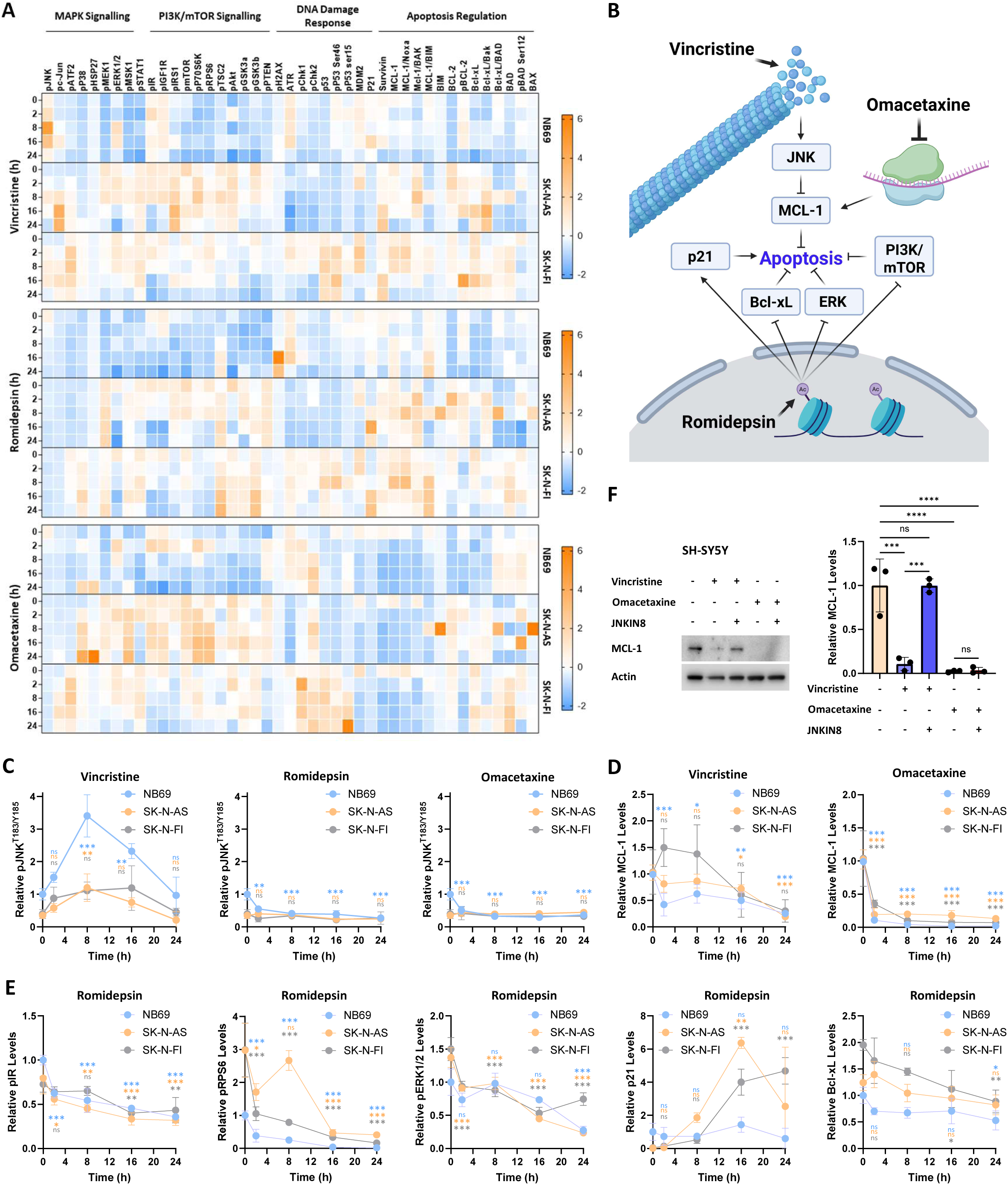
Mechanistic analysis of JNK-dependent and JNK-independent drugs. (**A**) Multiplexed analysis of signalling pathways and apoptotic regulators in NB69, SK-N-AS and SK-N-FI cell lines treated with either vincristine, romidepsin or omacetaxine (100 nM), at the time points indicated. Log2 transformed data is normalised to the median untreated value for each analyte across all cell lines (Mean, n=3). (**B**) A schematic of the mechanism of action for each individual drug. (**C**) Raw data for JNK phosphorylation (T183/Y185) from A, normalised to the zero time point for the NB69 cell line (Mean ± SD, n=3). (**D**) Raw data for MCL-1 expression from A, normalised to the zero time point for the NB69 cell line (Mean ± SD, n=3). (**E**) Raw data for the analytes indicated for the Romidepsin treatment arm in A. Data is normalised to the zero time point for the NB69 cell line (Mean ± SD, n=3). (**F**) Western blot analysis with the antibodies indicated in the SH-SY5Y cell line following treatment with vincristine (300 nM), omacetaxine (300 nM) or JNK-IN-8 (10 µM), as indicated (Mean ± SD, n=3). *** p<0.001, ** p<0.01, * p <0.05.

Multiple pathways have been implicated in the induction of apoptosis following treatment with romidepsin across diverse different tumour types (31). Interestingly, the activation of JNK-dependent apoptosis by romidepsin has previously been observed in malignant T-cells and hepatocellular carcinoma cells (32, 33), suggesting that the ability of this HDACi to activate JNK is highly cell-type specific. Other proposed mechanisms have included a decreased activation of PI3K/mTOR (32) and ERK (33) signalling, along with the accumulation of p21 (24, 34–37). Interestingly, alterations in all of these pathways were observed following romidepsin treatment, including a decrease in pIR, pRPS6 and pERK1/2, along with an increase in p21 levels in all but the NB69 cell line (**Fig. 3E**). While no consistent degradation of MCL-1 was observed, a significant decrease in an alternative Bcl-2 family member, Bcl-xL, was observed in the NB69 and SK-N-FI lines after 24 h of treatment (**Fig. 3B,E**).

In contrast, omacetaxine treatment resulted in the rapid degradation of MCL-1, with levels reaching <20% of pre-treatment baseline within 2 hours (**Fig. 3D**). This is in line with the role of omacetaxine as a protein synthesis inhibitor, and has been observed within other tumour types (38). These findings were also validated by Western blotting in the SH-SY5Y cell line (**Fig. 3F**), where a much slower, JNK-dependent loss of MCL-1 could be observed following vincristine treatment, whereas omacetaxine treatment resulted in a rapid, JNK-independent loss of MCL-1.

### Synergy between JNK-dependent and independent drugs

This convergence of both JNK-dependent vincristine and JNK-independent omacetaxine onto MCL-1 degradation has potential implications for understanding the response to pair-wise combinations of these, and other drugs. We have previously demonstrated that the standard-of-care drugs vincristine, doxorubicin and topotecan can all promote the degradation of MCL-1 (19). We therefore undertook synergy assays combining these JNK-dependent chemotherapy agents with our JNK-independent drugs, using both JNK-impaired (SK-N-AS, SK-N-FI) and JNK activating (SH-SY5Y) cell lines (**Fig. 4A,C**). For these assays, drug synergy was determined using the ZIP synergy score (**Fig. 4B,D**), revealing that the combination of either doxorubicin or topotecan with romidepsin was highly synergistic across all three lines (ZIP > 10), with synergy for the combination of vincristine and romidepsin only observed in the SK-N-FI cell lines (**Fig. 4B**). Conversely, all combinations between these standard-of-care drugs and omacetaxine resulted in either mildly antagonistic or additive drug interactions (ZIP <10) (**Fig. 4D**).

**Figure 4:**
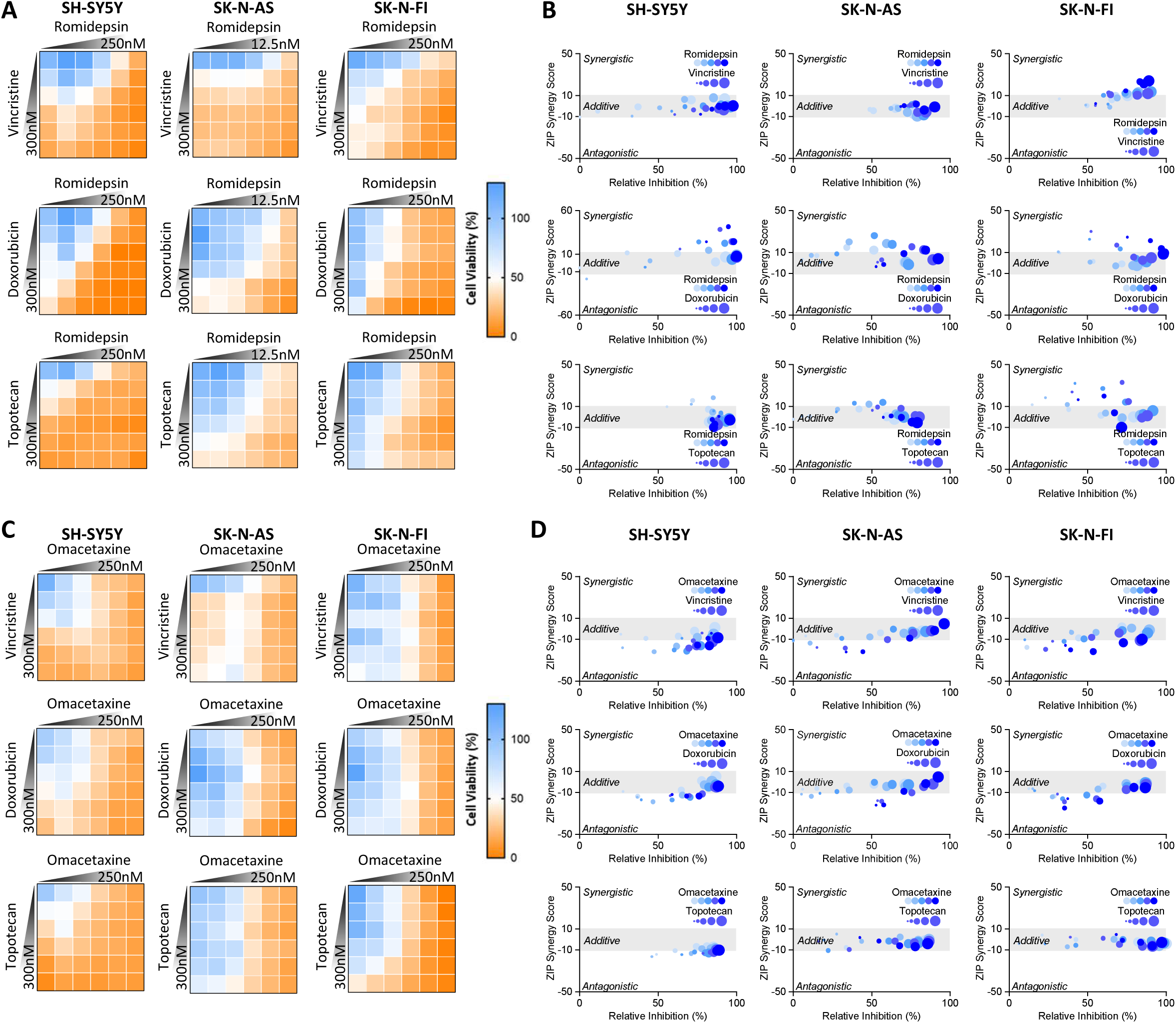
Synergy between JNK-independent drugs and standard-of-care chemotherapy. (**A**) Assessment of synergy performed using MTS assays in the SH-SY5Y, SK-N-AS and SK-N-FI cell lines with pair-wise combinations of vincristine, doxorubicin or topotecan with romidepsin. Dose responses were performed with a 1:2 dilution from the top drug concentration indicated (Mean, n=6). (**B**) Quantification of ZIP synergy scores from the data in A was performed with SynergyFinder 2.0. Values lower than −10 represent and antagonistic interaction, values higher than 10 a synergistic interaction. Increasing doses of standard-of-care chemotherapy are represented by are represented by an increase in size, while increasing doses of romidepsin are indicated darker shades of blue for each data point. (**D**,**E**) An identical set of cytotoxicity assays and synergy calculations performed with omacetaxine in the place of romidepsin.

While omacetaxine was identified as a JNK-independent drug, the lack of synergy between omacetaxine and other JNK-dependent chemotherapy agents may be due to their downstream convergence upon MCL-1 degradation for the induction of apoptosis. In contrast, romidepsin appears to act by a number of potential mechanisms to drive this cytotoxic response. Together, these findings increase the complexity of the task of identifying synergistic drug combinations, suggesting that a broader understanding of the network of apoptotic components involved in apoptotic signalling may be required to fully understand the factors underlying the emergence of synergy.

### Functional genomics analysis of drug synergy

Building upon these observations, we sought to perform a systems analysis of apoptotic signalling in response to a series of pair-wise drug combinations. To do this, we used our high-content readout of caspase 3/7 cleavage to quantitate drug synergy at the level of apoptosis induction and paired this data with a functional genomics screen using siRNA against 196 apoptosis network components (**Table S2**). This analysis was performed with a drug panel consisting of the JNK-dependent, standard-of-care drugs topotecan, SN38 (the active metabolite of irinotecan), vincristine and doxorubicin, as well as the JNK-independent drugs romidepsin and alisertib. Omacetaxine was not selected for this panel due to its lack of synergy in previous assays, while the less effective, but JNK-independent drug alisertib was chosen due to an emerging body of literature supporting the targeting of the protein kinase Aurora-A in neuroblastoma (39) and Phase I trials in combination with standard-of-care schedules (40).

Firstly, a direct readout of apoptosis was obtained for all possible pair-wise combinations within this drug panel in SH-SY5Y cells (**Fig. 5A**). This was performed using dosages that elicited an apoptotic readout of less than 50% at the maximum single-agent concentration for each drug, allowing potential synergistic responses to be observed. Drug synergy calculations were performed using the ZIP synergy score, once again revealing that romidepsin was highly synergistic in combination with all other drugs (ZIP > 10), with the most synergy observed in combination with topoisomerase inhibitors, SN38, topotecan and doxorubicin (**Fig. 5B**). This was not the case for other drug combinations, where very few synergistic combinations were observed.

**Figure 5:**
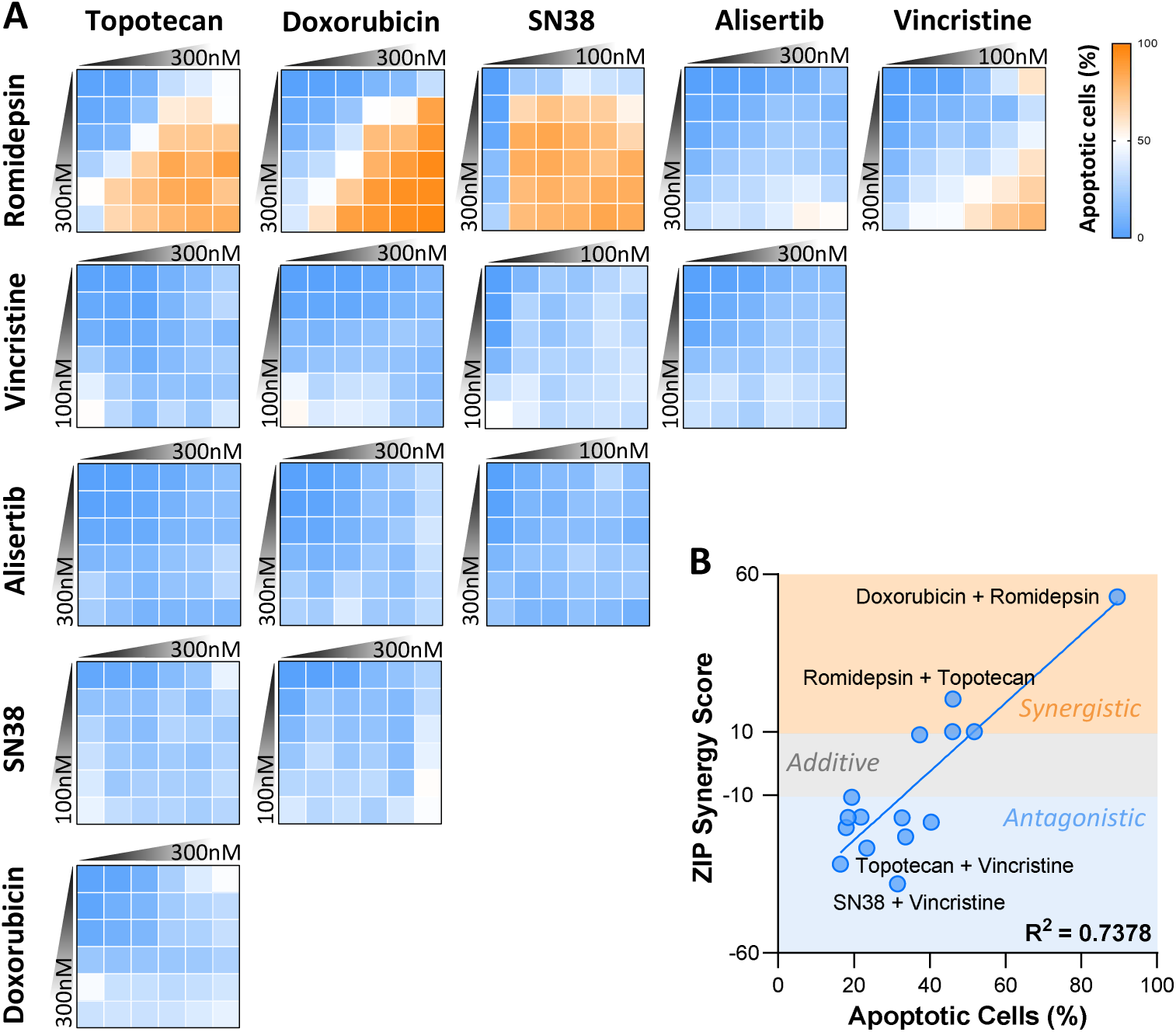
Apoptotic synergy assays. (**A**) Assessment of synergy based upon the percentage of cells with caspase activity after staining with NucView 488 (MFI > 20), measured by high content imaging in the SH-SY5Y cell line. Pair-wise dose responses were performed with a 1:2 dilution from the top drug concentration indicated (Mean, n=3). (**B**) Quantification of ZIP synergy scores from the data in A was performed with SynergyFinder 2.0. Values lower than −10 represent and antagonistic interaction, values higher than 10 a synergistic interaction. Increasing doses of drugs on the vertical axis are indicated darker shades of blue, while increasing doses for drugs on the horizontal axis are represented by an increase in size for each data point.

To further explore how this drug synergy emerges from the utilisation of apoptotic network components, a functional genomics screen was performed with an siRNA library spanning 196 apoptosis related proteins, followed by treatment with each of the individual drugs at their respective IC_30_ values and a high-content readout of apoptosis (**Fig. 6A, Table S2**). From this analysis, individual genes could be classified as either ‘essential’ for the induction of apoptosis (e.g. *CASP3*), or alternatively as ‘sensitising’ genes that resulted in increased chemotherapy-induced apoptosis following siRNA-mediated knockdown (e.g. *BCL2L1* – Bcl-xL) (**Fig. 6A, S2A**). Raw data from this screen was transformed and normalised, and quality control metrics were calculated (**Fig. S3A-E**). Confirming the fidelity of this dataset, unsupervised hierarchical clustering resulted in two small gene groupings containing additional genes with established roles as either universally essential (e.g. *CASP2*, *BBC3*) or sensitising (e.g. *PRKCD*, *STAT3*) across all drugs within the panel (**Fig. 6B**). In addition to this, two larger groups were also present, containing genes that were either broadly sensitising or essential, although with distinct drug specific patterns.

**Figure 6:**
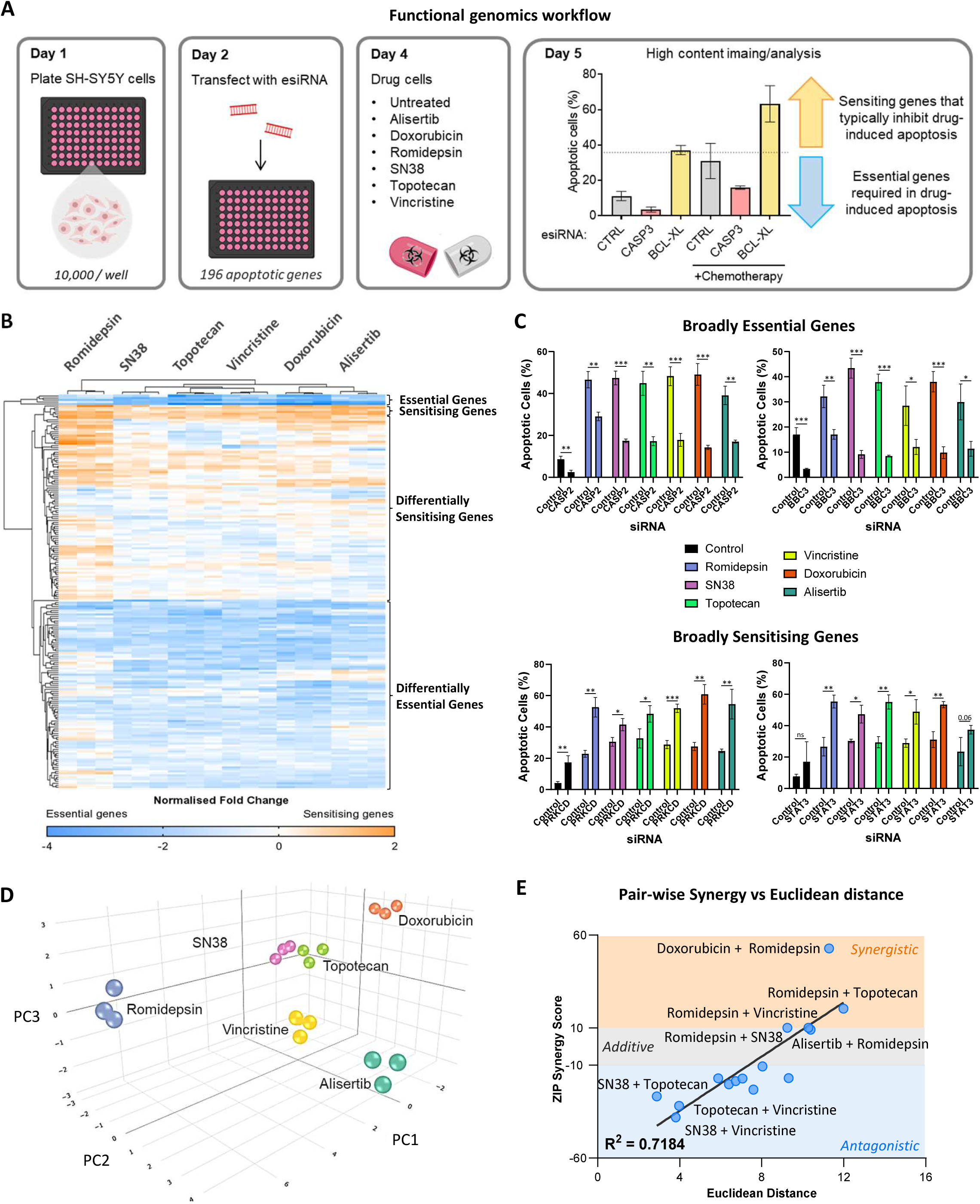
Apoptosis-focused functional genomics screen. (**A**) Schematic of the workflow for the functional genomics assay combining transfection of an esiRNA library of 196 apoptosis regulating genes followed by treatment with alisertib (50 nM), doxorubicin (50 nM), romidepsin (30nM), SN38 (15 nM), topotecan (50 nM) or vincristine (15 nM) and a high-content imaging based readout of apoptosis after staining with NucView 488 (MFI > 20). (**B**) Hierarchical clustering of log2 transformed screen data, normalised to the internal negative control for each plate. (**C**) Raw, plate level data for the individual genes indicated (Mean ± SD, n=3). (**D**) Three-dimensional XY plot of the first three components of a principal component analysis performed on screen level data. (**E**) Correlation between the Euclidean distance separating each pair-wise drug combination within the three-dimensional PCA plot in E and the respective ZIP synergy score from Figure 5B. *** p<0.001, ** p<0.01, * p <0.05.

Within this data, unexpected differences in the clustering of drugs were also apparent. While the related topoisomerase I inhibitors, SN38 and topotecan, clustered closely together, the unrelated microtubule inhibitor vincristine clustered closer to these drugs than did the topoisomerase II inhibitor doxorubicin, which clustered more closely with the Aurora-A kinase inhibitor alisertib. Romidepsin was a distinct outlier from the other drugs in this panel and appeared to induce a distinctly non-canonical form of apoptosis. Notably, romidepsin induced caspase activation that was independent of classical pro-apoptotic genes *PARP1* and *BAX* (**Fig. S2B**), as well as not relying upon expression of members of the p53 pathway (*TP53*, *TP53BP1*, *TP63*) (**Fig. S2C**), despite promoting the increased expression of p21 in previous experiments (**Fig. 3D**). Romidepsin also displayed a unique dependence on the pro-apoptotic *DIABLO* and the NF-kappa-B subunit *RELA*, which were not required for the induction of apoptosis by any of the other drugs in the panel (**Fig. S3D**). While knockdown of the anti-apoptotic *HMOX1* strongly sensitised the cells to romidepsin induced apoptosis, with only a modest impact upon the effect of doxorubicin (**Fig. S3D**).

Given this drug-dependent complexity, a principal component analysis was performed to better comprehend these emerging patterns and their relationship with drug synergy, which reduced this dataset to 17 components, of which the first 4 accounted for 75% of the variance (**Fig. S3F**). Analysis of the first two principal components, capturing 55.4% of the variance, confirmed the distinction between romidepsin and alisertib and the other drugs, while topotecan, SN38 and vincristine also clustered closely together (**Fig. S3G**). The third component chiefly provided a distinction between alisertib and doxorubicin (**Fig. S3H**), therefore the first three components were visualised together (65.7% of variance) (**Fig. 6D**). By rendering this data within a three-dimensional space, a clear distinction between romidepsin and the other drugs could be observed, again highlighting the distinct mechanism through which romidepsin activates apoptosis. Additionally, the close grouping of SN38 and topotecan with vincristine was also apparent. The mechanistic association of SN38 and topotecan would be expected, as both are topoisomerase I inhibitors and would likely activate apoptosis through similar mechanisms. However, as vincristine is a microtubule destabilising drug, this clustering was unexpected and suggested that while these drugs have different mechanistic targets, they ultimately activate apoptosis through very similar pathways.

Given each axis of this PCA plot represents an eigenvector grouping genes with a similar impact upon chemotherapy-induced apoptosis, distance within this multidimensional space was a direct result of differential utilization of these apoptotic network components by each drug. Therefore, to further investigate how these differences in apoptotic component utilisation impacted drug synergy, Euclidean distance within this three-dimensional space was measured for each pair-wise drug combination (**Fig. 6E**). These values were then plotted against the previously derived synergy data for each drug combination (**Fig. 5B**), revealing a good correlation (R^2^ = 0.7184) between these two parameters. Intriguingly, this analysis suggested that synergy between these drugs is associated with the magnitude of difference between the apoptotic signalling that they each induce as single agents, irrespective of their individual direct mechanistic drug target.

This multi-dimensional analysis again highlighted the finding that all romidepsin containing combinations were highly synergistic. In particular, ‘romidepsin + doxorubicin’ and ‘romidepsin + topotecan’ were the most synergistic combinations within this dataset. Conversely, all combinations between the similarly acting SN38, topotecan and vincristine were mildly antagonistic. In the clinical setting, combinations of these drugs form the backbone of many salvage therapy schedules for the treatment of relapsed or refractory HRNB, strongly suggesting that these existing multi-agent combinations may not be optimal and that the substitution of romidepsin within these regimens may provide greater clinical efficacy than the existing standard-of-care combinations.

### Implications for standard-of-care chemotherapy

Our previous studies with vorinostat revealed that this specific HDAC inhibitor was unable to sensitise patient-derived models of JNK-impaired, relapsed neuroblastoma to standard-of-care chemotherapy drugs within both *ex vivo* and *in vivo* experiments (19). However, our identification of romidepsin as a JNK-independent drug suggests that combining with this alternative HDAC inhibitor with existing standard-of-care drugs will be able to overcome this apoptotic impairment. Therefore, to demonstrate the efficacy of romidepsin within this JNK-impaired setting, we firstly utilised *ex vivo* cultures of these patient-derived xenografts, which were directly established with relapsed tumours from patients enrolled in the SIOPEN HR NB-1 clinical trial (41). These matched diagnosis and relapse models include CCI-NB01-DMC and CCI-NB01-RMT, along with CCI-NB02-DMB and CCI-NB02-RPT (**Fig. 7A**). The patient from which the CCI-NB01 models were established did not respond completely to Rapid COJEC and subsequently received two cycles of TVD, yet still developed progressive disease. In contrast, the CCI-NB02 patient had a complete response to Rapid COJEC but relapsed followed prolonged treatment delays due to surgical complications.

**Figure 7:**
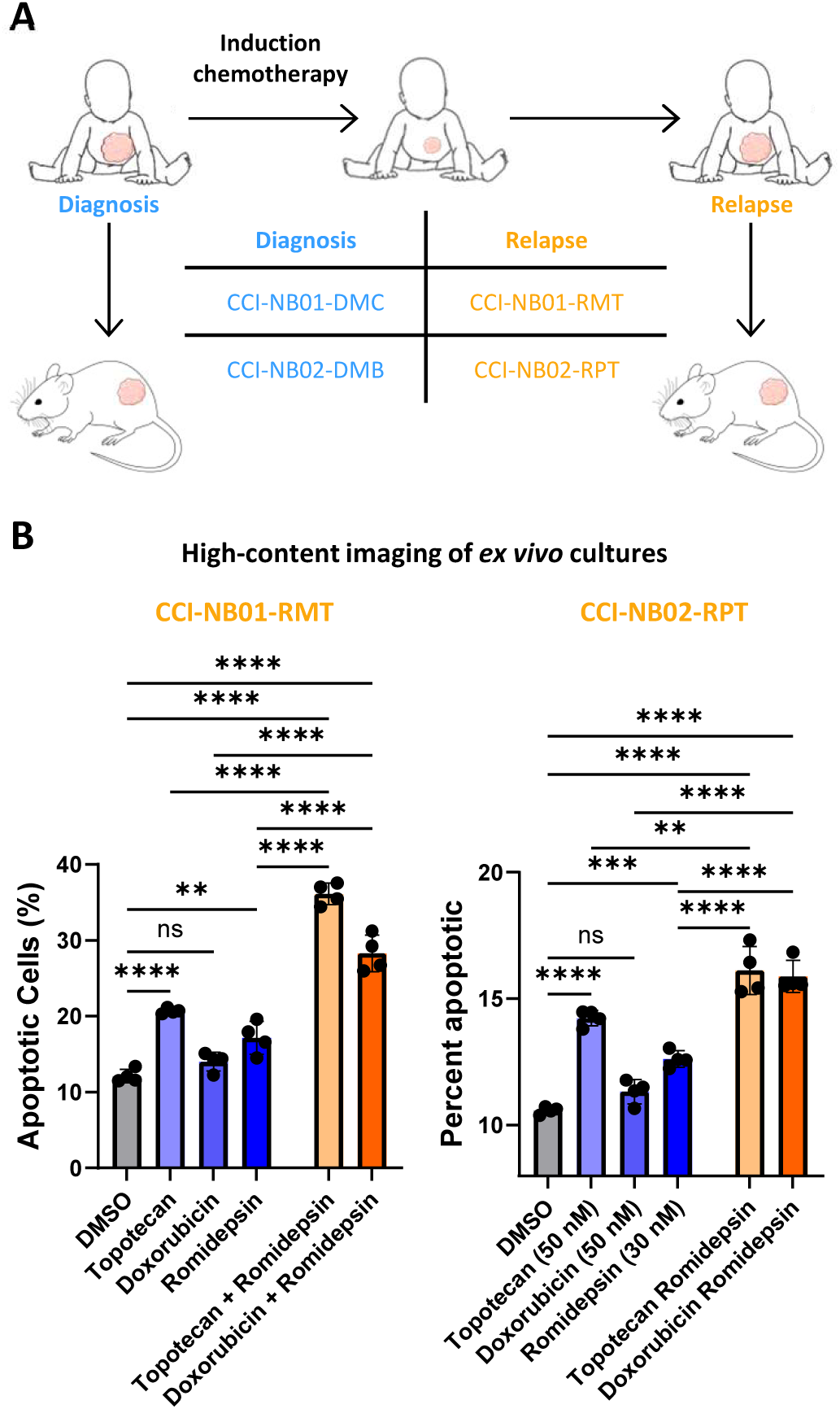
*ex vivo* cultures of relapsed PDX models. (**A**) Schematic of the matched diagnosis and relapse neuroblastoma PDX models. (**B**) High-content imaging based apoptosis assays performed on *ex vivo* cultures of the CCI-NB01-RMT and CCI-NB02-RPT PDX models following treatment with either topotecan (50 nM), doxorubicin (50 nM), romidepsin (30 nM) or the combinations indicated, for 24 h. The percentage of cells with caspase activity was quantified after staining with NucView 488 (MFI > 20) (Mean ± SD, n=3).

*Ex vivo* cultures from these relapsed PDX models were treated with topotecan, doxorubicin and romidepsin as single agents, along with both combination of topotecan + romidepsin and doxorubicin + romidepsin, prior to high-content imaging of apoptosis induction (**Fig. 7B**). Quantification of the percentage of apoptotic cells within each model again demonstrated the ability of romidepsin to significantly increase the induction of apoptosis when combined with these standard-of-care drugs, confirming the activity of this JNK-independent drug within these clinically relevant models of relapsed neuroblastoma. Therefore, we sought to further investigate the potential for these drug combinations to elicit a synergistic response within *in vivo* models.

Using the CCI-NB02-RPT PDX model, we firstly performed a dose response of topotecan treatment (0, 0.25, 0.5 and 1 mg/kg) for five consecutive days (**Fig. 8A**). As expected, increasing doses of topotecan significantly reduced tumour growth and prolonged survival compared to vehicle, which saturated at 0.5 mg/kg with no significant benefit between 0.5 and 1 mg/kg of topotecan (**Fig. 8B,C, S4A**). In order to combine romidepsin with this dose-dependent treatment of topotecan, a single dose of 1 mg/kg was used for this HDAC inhibitor. While this is lower than previously studies (42–44), dose-limiting toxicities were observed upon combination with topotecan at higher doses. Notably, at this low dose of romidepsin there was no single-agent impact upon tumour growth or survival (**Fig. 8B,C**). However, the benefit of romidepsin could still be seen in the reduction in tumour growth and significantly increased survival time when combined with the 0.25 mg/kg dose of topotecan. Median survival time increased from 26 days for single-agent topotecan to 31 days with the addition of romidepsin, rendering this combination as effective as the highest, single-agent dose of topotecan (1 mg/kg – 30 days median survival). This beneficial effect was also apparent in an analysis of tumour size at day 20, a time point at which the majority of vehicle and romidepsin tumours reached endpoint (**Fig. 8D**). Here, the combination of romidepsin with low dose topotecan (0.25 mg/kg) significantly reduced tumour growth compared to topotecan alone, and was as effective as the highest dose of 1mg/kg topotecan. Through the use of CombPDX (45) and InVivoSyn (46), this analysis confirmed the presence of synergistic interactions within this *in vivo* model, where the combination of romidepsin with both 0.25 and 0.5 mg/kg of topotecan gave a significant Bliss Synergy Score and the combination with 0.25 mg/kg a significant Global Bliss Combination Index (**Fig. 8E**)

**Figure 8:**
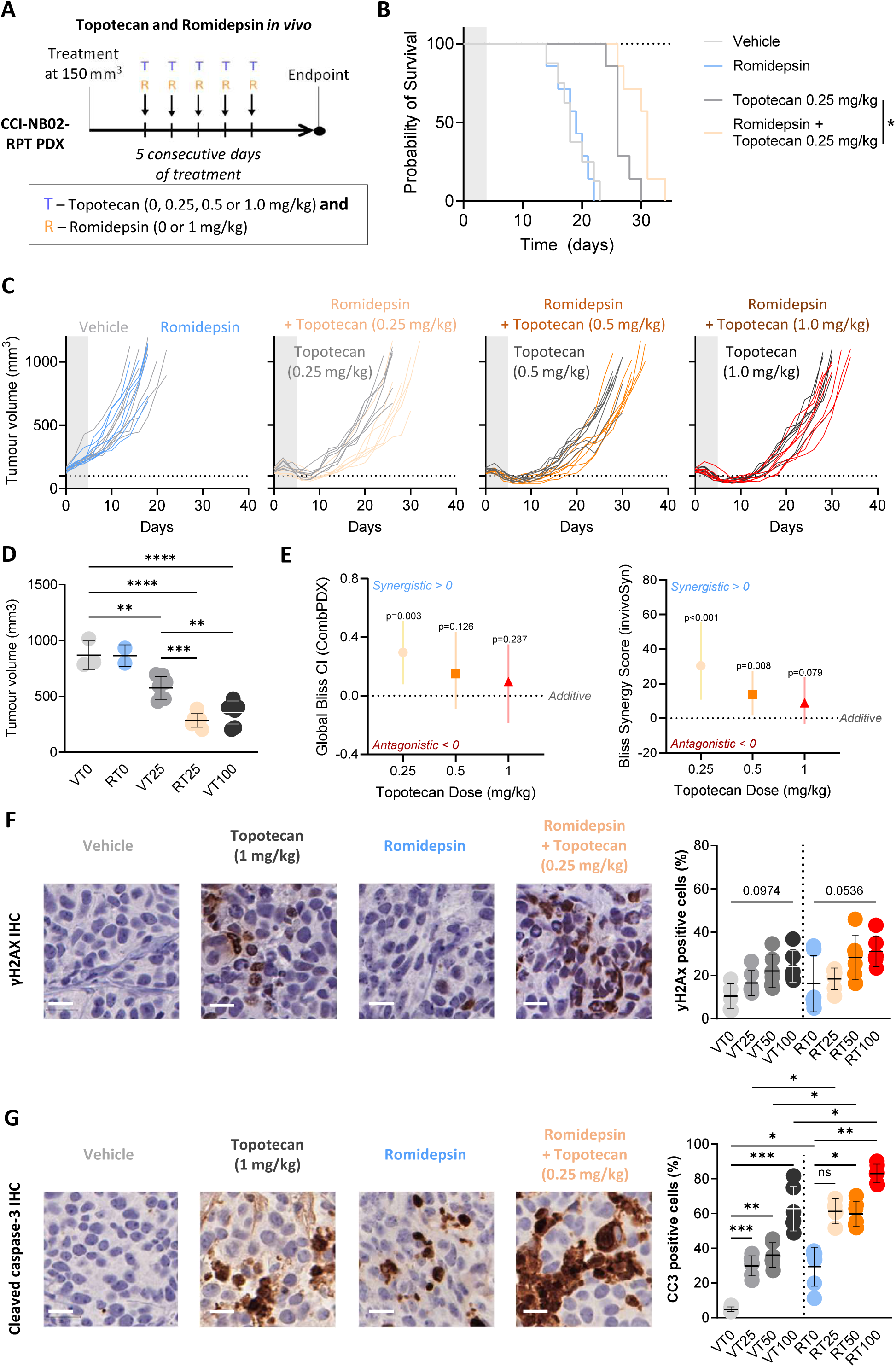
Synergy between romidepsin and topotecan *in vivo*. (**A**) Outline of the treatment schedule of the CCI-NB02-RPT PDX model with romidepsin in combination with increasing doses of topotecan, as indicated. NSG mice were implanted with 1×10^6^ CCI-NB01-RPT cells, once tumours reached 150 mm^3^ the mice received intraperitoneal injections of the drug combinations indicated, or the relevant vehicle controls, once daily for 5 days. Tumour growth was measured every day until ethical endpoint (1000 mm3) (**B**) Survival analysis for the treatment arms indicated. (**C**) Tumour growth curves (n=8). (**D**) Analysis of tumour volume at day 20 for the treatment arms indicated (**E**) *In vivo* drug synergy calculations using CombPDX (Bliss CI, bootstrap n=1000, CI=95%) and InvivoSyn (Bliss synergy score, bootstrap n=1000, CI=95%). (**F**) Immuno-histochemical analysis of γH2A.X staining. Representative images are shown for vehicle treatment (light grey), topotecan 0.25 mg/kg (grey), romidepsin alone (blue) and the combination of these two treatments (orange). Quantification is performed from staining performed on two separate sections, from three independent tumours (Mean ± SD, n=6). (**G**) Immuno-histochemical analysis of cleaved caspase 3 staining (Mean ± SD, n=6). All scale bars are 20 µm. *** p<0.001, ** p<0.01, * p <0.05.

This pattern was also recapitulated in the analysis of molecular markers by immunohistochemistry, performed on tumour samples collected following 2 days of treatment. Using γH2A.X staining as a marker of DNA damage induced by topotecan treatment, we observed a dose dependent increase in γH2AX positive cells, although this did not reach significance (**Fig. 8F**). While there was no significant difference in the induction of γH2AX positive cells with the addition of romidepsin, this was not the case for the induction of apoptosis, which was assessed by staining with a cleaved caspase-3 antibody (**Fig. 8G**). For this analysis, there was a significant, dose-dependent increase in apoptotic cells with single-agent topotecan treatment. However, upon combining this treatment with romidepsin, there was a further, significant increase in apoptosis induction at all topotecan doses.

### Including romidepsin within multi-agent chemotherapy schedules

While this analysis demonstrates the potential for romidepsin to synergise with these standard-of-care chemotherapies, these drugs are not used in isolation. Instead they are components of multi-agent schedules. Typical salvage regimens used for relapse and refractory HRNB currently include TOTEM (topotecan and temozolomide) and IT/TEMIRI (irinotecan and temozolomide). While these combination treatments do already provide a benefit for some patients, they are generally well tolerated and therefore often used as a backbone on which to test additional agents (13–15, 47–50).

While temozolomide was not effective within our initial drug screen (**Fig. 1E**), even at doses up to 3 µM, the combination of irinotecan (SN38) and romidepsin was identified as synergistic within our functional genomics analysis (**Fig. 5/6**) and represents another potentially clinically applicable combination. Therefore, romidepsin was assessed in combination with both TOTEM (rTOTEM) and IT (rIT), in the CCI-NB02-RPT relapsed PDX model (**Fig. 9A**). Individually, both TOTEM and IT reduced tumour growth and significantly improved survival compared to both vehicle and romidepsin alone (**Fig. 9B,C**). However, the inclusion of temozolomide within the TOTEM schedule did not actually improve upon the single-agent response of topotecan within our previous experiment (**Fig. S4A**), with a median survival of 29 days for single agent topotecan and 30 days for the TOTEM combination across these two experiments. While median survival increased to 34 days with the addition of romidepsin (rTOTEM), this did not reach significance. However, there was a significant improvement in survival for rIT compared to IT, with an increase in median survival from 39 to 48 days (**Fig. 9B,C**). Again, this improvement was evident despite the low dose of romidepsin used, which potentially limited the response in this context.

**Figure 9:**
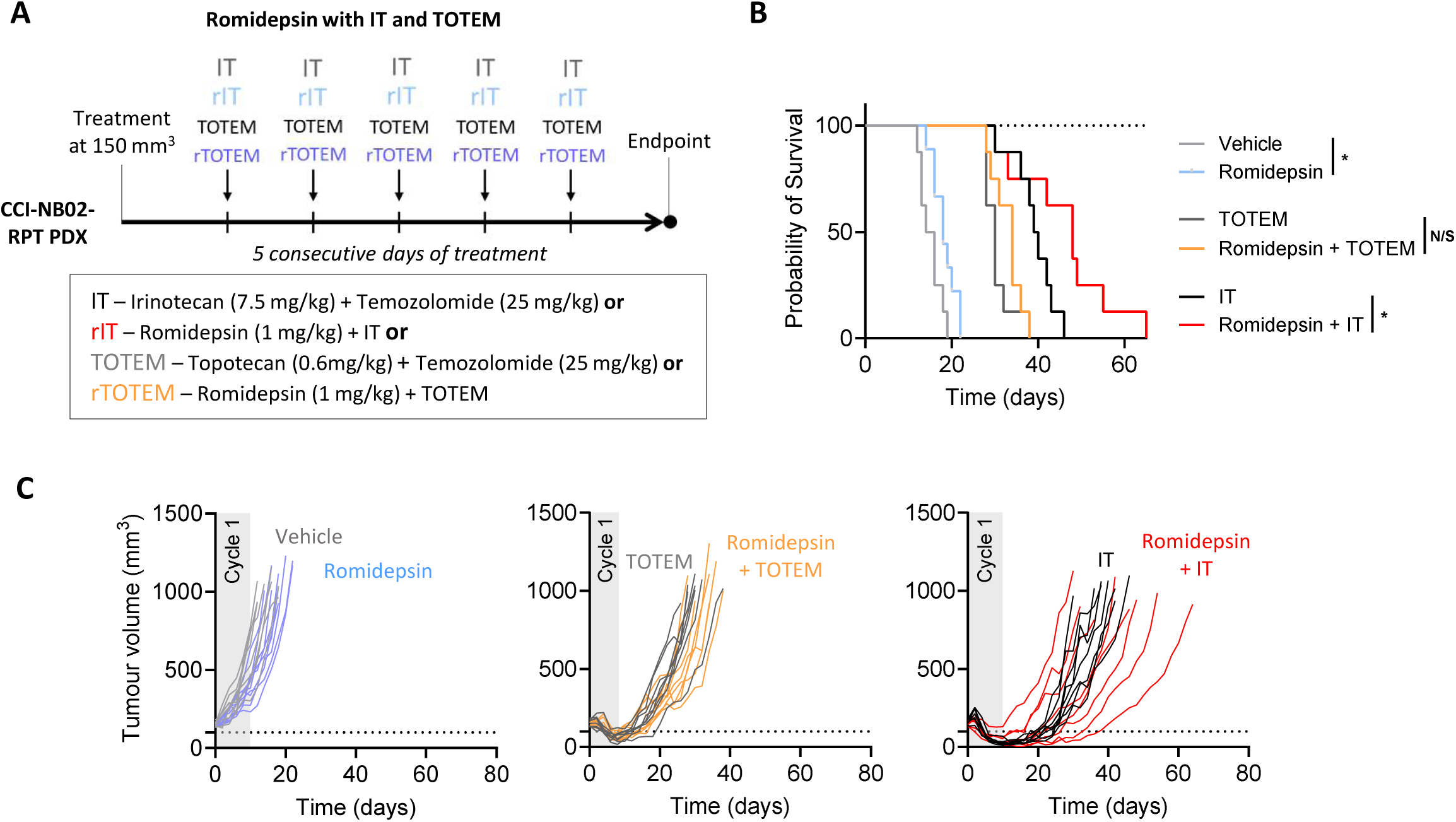
Combining romidepsin with standard-of-care backbones. (**A**) Outline of the treatment schedule of the CCI-NB02-RPT PDX model with romidepsin in combination with IT (Irinotecan + Temozolomide) and TOTEM (Topotecan + Temozolomide) as indicated. NSG mice were implanted with 1×10^6^ CCI-NB01-RPT cells, once tumours reached 150 mm^3^ the mice received intraperitoneal injections of the drug combinations indicated, or the relevant vehicle controls, once daily for 5 days. Tumour growth was measured every day until ethical endpoint (1000 mm^3^) (**B**) Survival analysis for all treatment arms. (**C**) Tumour growth curves (n=8). *p<0.05

Another candidate for investigation in this context is TVD salvage therapy, a 7 day protocol in which patients receive 5 days of topotecan followed by a 1 hour break, and then a 48 h continuous infusion of vincristine and doxorubicin. Based upon this timing of clinical drug administration, and the most significant pair-wise combinations within our dataset, we initially investigated whether topotecan-romidepsin-doxorubicin (TRD) and topotecan-vincristine-romidepsin (TVR) would be more effective than the existing TVD triple-agent combination. These triple-agent combinations were initially evaluated by simultaneous treatment within *in vitro*, high-content imaging apoptosis assays, optimised with single-agent concentrations of topotecan (50 nM), doxorubicin (30 nM), vincristine (15 nM) and romidepsin (30 nM) that induced a small (<5%) and non-significant increase in the levels of apoptosis in SH-SY5Y cells (**Fig. 10A**). Strikingly, within this assay, the triple-agent combination of TVD was unable to induce a level of apoptosis significantly higher than any of its single agent components, or the untreated control. However, both romidepsin containing triple-agent combinations (TVR and TRD) were capable of inducing significantly greater levels of apoptosis when compared with the standard-of-care TVD. This pattern of apoptosis induction was also apparent within all other cell lines included within the original cell line panel, including the JNK-impaired SK-N-AS and SK-N-FI lines, and also within the murine NB9464 neuroblastoma cell line (**Fig. 10A**). Colony forming assays were also conducted as an additional readout of drug response. In both the SH-SY5Y and NB9464 cell lines, colony formation was significantly impaired by all triple-agent combinations, although TVR and TRD were significantly more effective than TVD (**Fig. 10B**).

**Figure 10:**
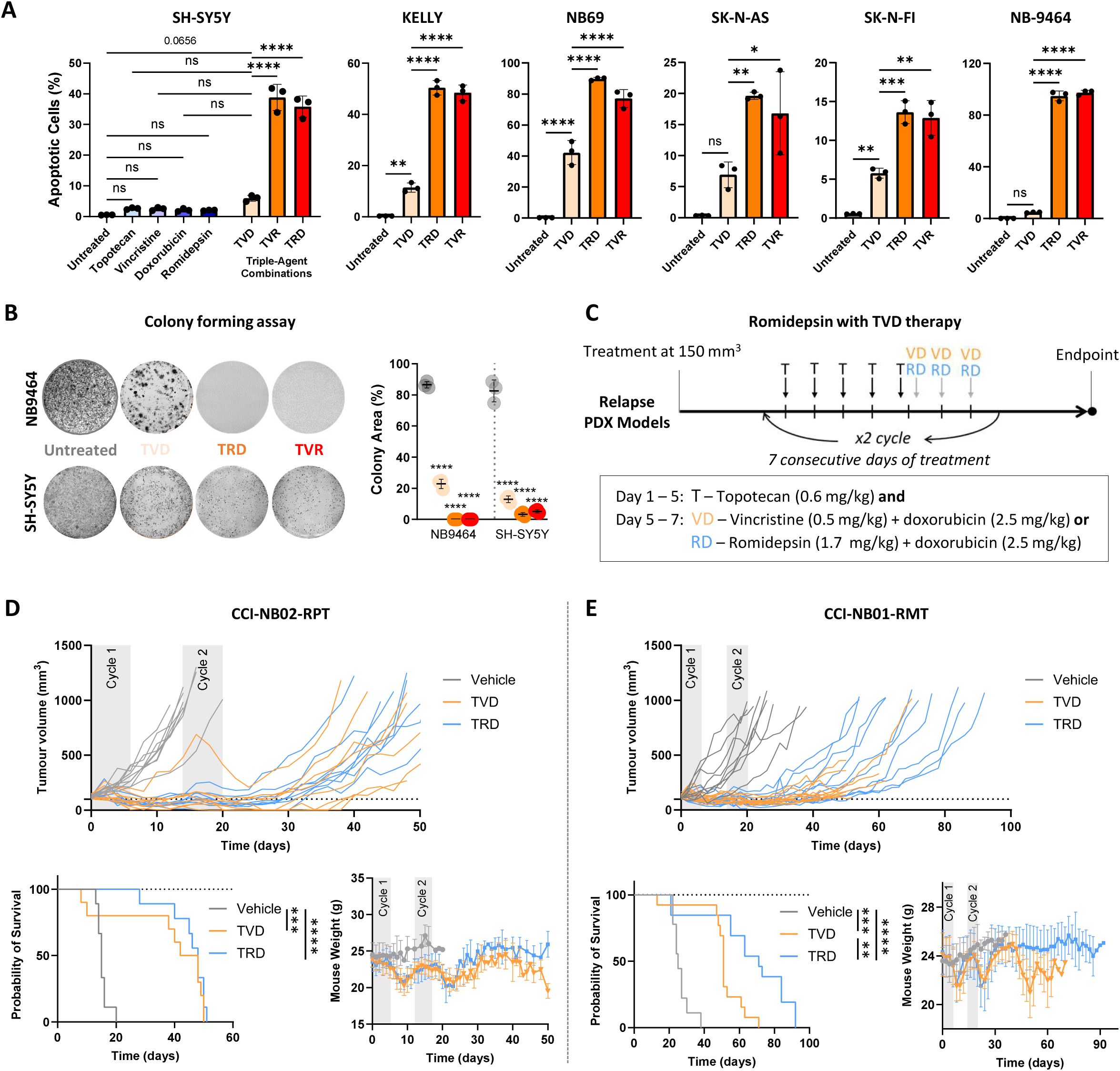
Inclusion of romidepsin within TVD. (**A**) High content imaging analysis of a panel of neuroblastoma cell lines after staining with NucView 488, following treatment with the triple-agent combination of topotecan (50 nM), vincristine (15 nM) and doxorubicin (50 nM) (TVD), or with romidepsin (30 nM) in the place of either doxorubicin (TVR) or vincristine (TRD). The percentage of cells with caspase activity was quantified after staining with NucView 488 (MFI > 20) (Mean ± SD, n=3). (**B**) Colony forming assays in NB-9464 and SH-SY5Y cells following treatment with TVD, TVR or TRD for 7 days (Mean ± SD, n=3). (**C**) Outline of the treatment schedule of the CCI-NB02-RPT and CCI-NB01-RMT PDX models with either TVD or TRD as indicated. NSG mice were implanted with 1×10^6^ PDX cells, once tumours reached 150 mm^3^ the mice received injections of the drug combinations indicated (IP – topotecan, romidepsin; IV – vincristine, doxorubicin) or the relevant vehicle controls, at the timing indicated. (**D**) Tumour growth was measured every day until ethical endpoint (1000 mm^3^) (Mean ± SD, n=8), upon which the survival analysis was based. Mouse weights were also measured throughout the course of treatment and tumour growth (Mean ± SD, n=8). **** p<0.0001, *** p<0.001, ** p<0.01, * p <0.05.

Given the potential to evaluate romidepsin in a triple-agent combination with its two most synergistic pair-wise drug partners, we further investigated the TRD treatment schedule *in vivo* using both the CCI-NB02-RPT and CCI-NB01-RMT PDX models, as this latter model was derived from a patient that had relapsed following treatment with TVD. For these studies, subcutaneous tumours were treated with two cycles of a schedule designed to mimic either the standard-of-care TVD (topotecan 0.6 mg/kg, IP; vincristine 0.5 mg/kg IV; doxorubicin 2.5 mg/kg) or the modified TRD schedule (topotecan 0.6 mg/kg, IP; romidepsin 1.7 mg/kg, IP; doxorubicin 2.5 mg/kg, IV) (**Fig. 10C**).

Within both models, the TVD and TRD schedules impaired tumour growth and significantly increased survival compared to vehicle (**Fig. 10D,E**). Within the CCI-NB02-RPT model there was no benefit of TRD over TVD, although there was a significant improvement in survival for the modified TRD schedule apparent within the CCI-NB01-RMT PDX, where median survival increased from 51 to 71 days. In this model, late-onset toxicity resulting from TVD treatment was also evident from the rapid weight loss occurring from 50 days after treatment initiation, which did influence this survival difference (**Fig. 10E**). Notably, this weight loss did not occur in mice treated with TRD, suggesting that this modified schedule may be more tolerable than the existing standard-of-care.

## Discussion

Although significant focus has been placed upon genomic studies in HRNB, the development of a broadly applicable precision medicine approach has been hampered by the lack of frequently occurring, therapeutically actionable mutations. Outside of the emergence of ALK inhibitors and GD2 immunotherapy, multi-agent chemotherapy remains standard-of-care for this aggressive disease. Unlike targeted therapies, which are rationalised by the presence of specifically actionable mutations, these combination chemotherapy schedules have been historically assembled by selecting drugs with different mechanistic targets, in an attempt to increase anti-tumour activity and prevent the onset of resistance (16). However, our findings suggest that synergy between chemotherapy drugs does not necessarily emerge from the combination of drugs with different mechanisms of action, rather from drugs that differ in their utilisation of apoptotic network components.

This finding is most apparent in the combination of the microtubule inhibitor vincristine with either the topoisomerase I inhibitors topotecan/SN38, or the topoisomerase II inhibitor doxorubicin. While each of these chemotherapy drugs exert their action upon cancer cells through different direct targets and different mechanisms, they ultimately activated apoptosis in a similar manner within our functional genomics screen. As a result, these drug combinations produced weakly additive, or even antagonistic responses within our quantitative apoptosis assays. This data is even more striking when considering that these drugs – topotecan, vincristine and doxorubicin – are the components of the recently used, standard-of-care TVD schedule for relapsed/refractory neuroblastoma.

TVD salvage therapy was first used in a phase II trial for refractory and relapse advanced neuroblastoma in 2003, giving a combined response rate of 64% (51). However, TVD is no longer recommended for patients who do not respond to Rapid COJEC as there was no evidence of an improvement in overall or event free survival for these patients (52). Interestingly, our *in vitro* data suggested that there is little benefit from the TVD triple-agent combination in terms of apoptosis induction, although there was a robust and significant improvement upon the substitution of romidepsin into this combination. However, the benefit of including romidepsin within a modified TRD schedule was limited within our *in vivo* studies, likely due to both the low dose romidepsin required to circumvent toxicity and the timing of drug delivery, which involved an initial 5-day period of single-agent topotecan treatment before the introduction of the other components, reducing the potential for a synergistic response.

In line with the limitations of the TVD schedule, it is now being replaced by alternative combinations that involve concurrent treatment with temozolomide and either topotecan (TOTEM) or irinotecan (IT/TEMIRI). The use of temozolomide within these combinations was driven by its tolerability in phase I pediatric trials, ability to cross the blood-brain barrier and a distinct mode of action from these topoisomerase inhibitors (53). However, in our stratified drug screen, temozolomide was not capable of activating apoptosis within neuroblastoma cell lines and did not actually improve upon single-agent topotecan within our relapsed PDX model. Importantly, the benefit of combining romidepsin with topotecan could be seen in the synergistic *in vivo* response in this PDX model, along with the improvement in survival seen with the addition of romidepsin to IT. Despite the limitation of romidepsin dosage within these models, the observation of *in vivo* synergy with this HDAC inhibitor in these aggressive models of relapsed HRNB warrants further investigation to overcome these dosage limitations.

Despite numerous different drug combinations being used widely across many advanced cancers, only a fraction of these are considered truly synergistic (54). These synergistic drug combinations may not only improve response rates, but also reduce toxicity by allowing the use of lower doses of each individual drug. Therefore, significant, highly collaborative efforts have focused on the identification of synergistic drug combinations through high-throughput screening and correlation with molecular features, including gene expression signatures, mutation status and synthetic lethal interactions (55–59). While these approaches have demonstrated that the prediction of drug synergy is possible, overall probability is often limited, suggesting that a deeper understanding of how synergy emerges from drug combinations is needed. Our finding that the differential utilisation of apoptotic pathway components for each individual drug is a key indicator of pair-wise drug synergy has the potential to add an additional, mechanistic layer to these predictive approaches which may improve their performance.

Interestingly, mechanism-based studies focused on the direct perturbation of specific proteins and the use of targeted therapies have demonstrated that likelihood of observing a synergistic response increases when targeting proteins with either strong functional similarity or dissimilarity (60). This additional complexity may arise specifically from the combination of targeted therapies involving multiple components of the same signalling pathway. A key example of this is the combination of BRAF and MEK inhibitors, which display synergy by overcoming the negative feedback induced by single-agent BRAF inhibition (60). While our data suggests that targeting different apoptotic pathways simultaneously is beneficial in the context of combining chemotherapy drugs to overcome drug resistance, this may not always apply in the context of specific combinations of targeted therapies.

In pursuing the investigation of drug resistance in HRNB, our studies continually highlight the potential for HDAC inhibitors to improve the response to standard-of-care chemotherapies (18, 19). Indeed the central involvement of HDACs in the pathobiology of neuroblastoma has been highlighted by both a multitude of preclinical studies and the performance of clinical trials testing the HDAC inhibitors valproic acid, vorinostat and sodium phenylbutyrate in pediatric settings (61). However, none of these findings have advanced the use of HDAC inhibitors into clinical practice for HRNB. It is likely that this limitation lies more within the lack of HDAC specific drugs rather than an intrinsic flaw in the concept of targeting HDACs. Indeed, many studies have identified a strong discordance between the functional consequences of genetic inactivation/depletion of HDACs and treatment with HDAC inhibitors, while also attributing the function of many of these drugs to non-histone, off-target proteins (62, 63).

While there are well noted issues with specificity for HDAC inhibitors, there are also ongoing issues with toxicity that are likely on-target in nature. Indeed, a number of adverse effects have been reported in patients receiving romidepsin treatment in clinical trials for glioblastomas, renal cell carcinomas, colorectal cancer (64–66), which has limited its clinical implementation. We also observed dose-limiting toxicity in our mouse models with the use of romidepsin when combined directly with topotecan, which was otherwise predicted to be a highly synergistic drug combination. Therefore, the reduced dose required for treatment of our PDX models likely limited the efficacy within our studies, although we were still able to observe synergy emerging from this drug combination.

It is therefore very likely that future studies targeting HDACs in HRNB, and indeed other tumour types, will need to move away from these pan-HDAC inhibitors and begin to leverage new classes of drugs capable of providing alternative methods of drug delivery or target inhibition. This will likely include approaches such as antibody-drug conjugates containing HDAC inhibitor payloads, which may limit side effects by direct delivery of these drugs into tumour cells. While the degradation of specific HDACs through the use of PROTACs is also another approach potentially capable of providing the specificity required to target this complex family of enzymes.

Future studies will reveal whether our ability to target HDACs will improve with the emergence of these next-generation therapeutics. However, our findings provide a clear rationale to think differently about the design of effective multi-agent chemotherapy schedules for HRNB, and indeed other tumour types that still rely on chemotherapy as frontline treatment. While an understanding of the fundamental basis of drug synergy will be important, another key consideration for the treatment of relapsed tumours is consideration of the extensive rewiring that occurs within these apoptotic pathways following an initial exposure to chemotherapy (19). Because of this, many effective drugs, and drug combinations, identified using primary tumours models will not continue to be effective within the relapse setting, where the biology of the tumour has been fundamentally altered. A focus on the mechanisms that underlie these adaptive responses and their ability to influence drug synergy, along with studies designed to identify effective drug combinations in this specific context, will be essential if we are to improve outcomes for patients with this aggressive disease.

## Materials and Methods

### Study design

All *in vitro* experiments were performed and quantified in at least biological triplicates, with relevant sample sizes reported for each experiment. All single-cell, high content imaging experiments were performed with n>1000 for each treatment/condition. *In vivo* xenograft studies were randomized, with relevant group sizes reported for each individual experiment.

### Reagents

The source and details for all reagents are listed in the supplementary information, including antibodies and drugs (Table S3).

### Experimental Model and Subject Details

The Kelly, SH-SY5Y, NB69, SK-N-AS, SK-N-FI and IMR-32 neuroblastoma cell lines were all maintained in phenol-red RPMI-1640 media with 10% Foetal Calf Serum (FCS), and Penicillin/Streptomycin under standard tissue culture conditions (5% CO_2_, 20% O_2_). All cell lines were authenticated by short tandem repeat polymorphism, single-nucleotide polymorphism, and fingerprint analyses, passaged for less than 6 months, and confirmed as negative for mycoplasma contamination using the MycoAlert luminescence detection kit (Lonza, Switzerland).

The generation, maintenance, and propagation of the two sets of matched diagnosis and relapse models (CCI-NB01 and CCI-NB02) have been previously described (19, 41). All tumour specimens were obtained from patients at the Sydney Children’s Hospital Network (SCHN) under approval by the SCHN Human Research Ethics Committee (LNR/14/SCHN/392, LNR/14/SCHN/497). Informed parental or guardian consent was obtained for each patient.

### Western blotting

Cell lysates were washed twice with ice cold PBS, harvested using lysis buffer containing 50 mM Tris-HCl pH 7.4, 150mM NaCl, 1mM EDTA, 1 % (v/v) Triton X-100, 0.2 mM sodium orthovanadate and 1% (v/v) protease inhibitor cocktail, and kept on ice. Samples were centrifuged at 12,000 rpm at 4 °C, and the supernatant was collected. Protein concentration of lysate samples was performed with a Bradford assay and measured at 595nM with a FluoSTAR Omega plate reader. Lysates were diluted to 1 mg/mL with 1X (v/v) NuPAGE LDS sample buffer added and denatured at 95°C for 2 min. SDS-PAGE electrophoresis and western blotting was performed using the NuPAGE SDS PAGE Gel System and NuPAGE Bis Tris Precast Gels (4-12%) (Life Technologies). Western Lightning PLUS Enhanced Chemiluminescent Substrate (PerkinElmer) was for imaging western blots on the Vilber Lourmat Fusion chemiluminescent imaging system. Quantitative western blotting was performed using multistrip western blotting.

### Mathematical Modelling

All simulations of JNK activity were performed using a dynamic ODE model constructed through rule-based modelling that we have fully described previously (18). Briefly, this model consists mainly of the three-tiered JNK kinase cascade - a MAP3K (ZAK), two MAP2Ks (MKK4 and MKK7) and MAPKs (JNK1/2). In addition to this core structure, the model also contains a positive feedback loop mediated via JNK phosphorylation of Thr66 and Thr83 in the N-terminal region of MKK7, and inhibitory crosstalk through Akt-mediated phosphorylation of MKK4 (Ser80) and MKK7 (Thr385). All simulations and predictions were performed following the calibration and validation steps outlined in our previous work and executed using MATLAB R2020a.

To simulate JNK activity in different cell lines, input parameters were generated by measuring the relative abundance of each component of the model (I.e. ZAK, MKK7, MKK4, JNK and phosphorylated Akt Ser473) through quantitative western blotting and densitometry. As this model was initially constructed using the SH-SY5Y cell line, all values were normalised to those obtained for the parental SH-SY5Y cell line.

### Cell-based assays

#### Cytotoxicity assay

Cytotoxicity assays were performed as per manufacturer’s instructions for CellTiter 96 Aqueous proliferation assay, as previously described (67). In short, cells were plated at 1 × 10^5^ cells/well. Once adherent, cells were treated with individual or combination chemotherapy as indicated in each experiment. Following 24 h, CellTiter MTS reagent was added and incubated for 2 – 4 h. Absorbance was measured at 490 nm with a FluoSTAR Omega plate reader. Drug synergy was performed using SynergyFinder2 (68). The analysis of sub-G1 populations by propidium iodide staining and flow cytometry was performed as previously described (69).

#### Colony forming assay

Neuroblastoma cell lines were plated on collagen coated 12 well tissue culture plates at 5 × 10^5^ cells/well. After 24 h, triplicate wells were then treated with chemotherapy for 7 days before media was replenished with fresh culture media. Cells were allowed a 7 – 10 day recovery period before plates were stained. Briefly, 12 well plates were fixed with ice-cold methanol for 20 min at 4 °C. Cells were washed in water before being stained with a 0.1 % crystal violet solution as per manufacturer’s instructions. Excess crystal violet stain was washed off, then dried overnight and imaged using a scanner. Images were analysed using the ColonyArea ImageJ plugin (70).

### High-content imaging

High-content imaging analysis was performed with a Cellomics Arrayscan VTI HCS Reader. For all assays, imaging and parameter optimisation, cell gating and preliminary data analysis was conducted in the native HCS Studio 2.0 Cell Analysis Software (6.0.3.4020) and raw data was exported for further analysis.

#### Apoptosis Assay

Live cell imaging of apoptosis was performed with NucView 488 or 530 Caspase-3 enzyme substrate. In short, cells were seeded at 10,000 cells per well 100 μL of culture media in black 96-well plate and cultured overnight. Chemotherapy drugs were added to each well in 50 μL of culture media and incubated for 24 h. Prior to imaging, cells were stained with 1 μM of NucView 488 or 530 Caspase-3 enzyme substrate and 1 μg/mL Hoechst 33342 for 20 min at 37 °C. Nuclear staining with Hoechst 33342 was imaged on the 358/408 channel and a nuclear mask was rendered using the “CompartmentalAnalysis.V4” analysis template. Nuclear fluorescence intensity of the NucView 488 was imaged on the 485/524 channel and NucView 530 on the 549/593 channel. Mean fluorescent intensity (MFI) was measured and analysed as a direct readout of apoptotic, cleaved caspase 3/7 activity.

Cells with a cleaved caspase signal over a threshold of 20 MFI were identified as apoptotic and calculated as a percentage of the whole cell population as previously published (19).

#### Functional genomics screen

A functional genomics screen was carried out with a custom library of 196 × esiRNA of apoptosis network components (Table S2), with seven drug treatment arms (control, alisertib, doxorubicin, romidepsin, SN38, topotecan and vincristine). Firstly, 10,000 SH-SY5Y cells were plated in 100 μL of culture media in black 96 well plates. The "edge effect” was mitigated by only utilising the internal wells of a 96-well plate, while external wells were filled with 200 uL of PBS. Following 24 h, cells were transfected in triplicate, using jetPRIME. Transfection of the esiRNA library was performed in batches of 8 × esiRNA at a time, along with a Firely Luciferase (FLUC) esiRNA negative control and caspase-3 esiRNA positive control for each of the seven treatment arms. Each transfection mixture was made up with 300 ng of esiRNA (or 1.5 μL of 200 ng/μL esiRNA stock), 0.75 μL of jetPRIME and 50 μL jetPRIME buffer, which was dispensed at 10 μL/well. Each well contained 60 ng/μL of esiRNA, 0.15 μL of jetPRIME in 10 μL of jetPRIME buffer. Cells were stained with 1 μM of NucView 488 and 1 μg/mL Hoechst 33342 for 20 min at 37 °C, before imaging as described above.

### Fluorescence Microscopy of Antibody Staining

Cells for immunofluorescence imaging were plated on collagen coated tissue culture treated plasticware. Cells were fixed in 4 % PFA for 20 min, before being permeabilised in 0.1 % (v/v) Triton-X in PBS. Cells were blocked in 2 % (w/v) BSA in PBS blocking buffer for 1 h. Primary antibodies were diluted as per product specifications overnight in blocking buffer at 4 °C. Unless using a primary antibody conjugated to a fluorophore, cells were incubated with the respective secondary antibody 1:400 in blocking buffer for 1 h. Nuclei were stained with DAPI at a 1:1000 dilution in MilliQ water. Slides were imaged on the SP8 basic confocal microscope or scanned on the Cellomics Arrayscan VTI HCS Reader.

Immunofluorescence of protein synthesis was performed using Plus OPP Alexa Fluor™ 488 Protein Synthesis Assay Kit as per manufacturer’s instructions before being imaged on the Cellomics Arrayscan VTI HCS Reader.

### Multiplex analysis

Multiplex analysis was performed using a Bio-Plex MAGPIX system (#171015044) and Bio-Plex Pro-Wash Station (Bio-Rad) as previously described (71). Lysates were analysed using the Bio-Plex Pro RBM Apoptosis Panel 3 (Bio-Rad), the MILLIPLEX MAPK/SAPK Signaling Kit, the MILLIPLEX Akt/mTOR Phosphoprotein Kit, MILLIPLEX DNA Damage/Genotoxicity Kit, the MILLIPLEX MAP Bcl-2 Family Apoptosis Panel 1, and the MILLIPLEX MAP Bcl-2 Family Apoptosis Panel 2 (Merck).

### PDX models

#### In-vivo models

Following expansion of each PDX in NOD-SCID-gamma (NSG) mice, dissociated PDX tumour cells were subcutaneously engrafted into 6-week old secondary NSG mice (1×10^6^ tumour cells/graft in Matrigel) and randomised into treatment groups. Relevant treatment schedules for each drug or priming combination commenced once tumours reached 150 mm^3^. Topotecan (0.25 – 1 mg/kg), romidepsin (1 – 1.7 mg/kg) and irinotecan (7.5 mg/kg) were all delivered in a PBS vehicle by intraperitoneal injection. Temozolomide (25 mg/kg) was delivered in water by oral gavage. Vincristine (0.5 mg/kg) and doxorubicin (2.5 mg/kg) were delivered by intravenous injection in a PBS vehicle.

Tumour growth was measured every day and growth curves generated. Ethical endpoint was determined by ulceration, tumour size or maximum weight loss. Time to end-point was assessed using Kaplan-Meier survival curves, which were compared using the Log-rank test. All statistical analyses were performed using built-in functions in GraphPad Prism (Version 9, GraphPad Software).

#### Ex-vivo cultures

To generate *ex vivo* patient derived cell cultures from these PDX models, ∼20-30×10^6^ dissociated cells from each model were plated onto laminin-coated 15cm tissue culture dishes in IMDM media containing 20% FCS and Insulin/Transferrin/Selenium supplement (1:500). Following a 24 h incubation under standard tissue culture conditions (5% CO_2_, 20% O_2_), the tumour cells were harvested with Puck’s EDTA (140 mM NaCl, 5 mM KCl, 5.5 mM glucose, 4 mM NaHCO_3_, 13 µM Phenol Red, 0.8 mM EDTA, and 9 mM HEPES), which facilitated the selective detachment of tumour cells while stromal fibroblasts remained attached. Tumour cells were then processed for functional assays as indicated.

### Immunohistochemistry

Immunohistochemistry was performed on formalin-fixed paraffin-embedded sections using the Leica BOND RX. Slides were first dewaxed and rehydrated. Heat-induced antigen retrieval was performed with citrate (pH 6) retrieval buffer for 20 min at 100°C. Primary antibodies were diluted 1:500 (ki67), 1:200 cleaved caspase-3 and 1:2000 (γH2AX) in Leica antibody diluent and incubated for 60 min on slides. Antibody staining was completed using the Bond Polymer Refine immunohistochemistry protocol and reagents. Slides were counterstained on the Leica Autostainer XL. Bright-field images were taken on the Aperio CS2 Slide Scanner Quantification of single-cell staining intensity was performed using the cell detection function of QuPath (v0.2.3).

### Data Analysis

#### Statistical tests

Statistical tests were performed as required and specified. All data analysis and graphing of data was performed using GraphPad Prism unless otherwise specified.

#### Analysis of drug synergy

Drug synergy analysis was conducted using ZIP synergy calculations on SynergyFinder 2.0 (68). The ZIP synergy model was chosen as it combines the advantages of both the Loewe and Bliss models, allowing greater tolerance of experimental noise, which is common in large experimental screens and provides an improved solution for identifying true synergistic interactions, while keeping the false positive rate relatively low (72).

#### Functional genomics screen

Analysis of the esiRNA functional genomics screen was carried out in three parts using Perseus proteomic analysis software (73). Firstly, raw data was collected from HCI analysis as triplicate measurements of the percentage of cells undergoing apoptosis. Secondly, plate level data was normalised to the negative control (FLUC esiRNA) and collated into screen level data spanning the entire esiRNA library. Screen level data was log2 transformed to reflect fold changes in activation of apoptosis. Thirdly, systems analysis and quality control metrics were performed for the dataset. To assess the quality of the functional genomics screen data, a strictly standardized mean deviation (SSMD) score was calculated for screen level data per plate per treatment arm. The formula for SSMD is:

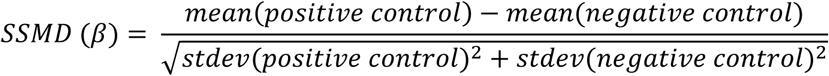

A SSMD takes into consideration the difference between the positive and negative control, and for RNA interference high throughput screening, a SSMD (β) around 0.5 is indicative of adequate data quality strictly standardized mean deviation or SSMD (β) as a metric for screen data quality. SiRNA screens tend to provide poor quality data with β ∼ 0.5 as minimum cut-off as follows (74, 75).

| Data quality | Criteria |  |  |  |
| --- | --- | --- | --- | --- |
|  | Moderate control | Fairly strong control | Strong control | Strong control |
| Excellent | $\beta \geq 2$ | $\beta \geq 3$ | $\beta \geq 4.7$ | $\beta \geq 6.67$ |
| Good | $2 > \beta \geq 1$ | $3 > \beta \geq 2$ | $4.7 > \beta \geq 3$ | $> \beta \geq 4.7$ |
| Inferior | $1 > \beta \geq 0.5$ | $2 > \beta \geq 1$ | $3 > \beta \geq 2$ | $4.7 > \beta \geq 3$ |
| Poor | $\beta < 0.5$ | $\beta < 1$ | $\beta < 2$ | $\beta < 3$ |

Unsupervised hierarchical clustering and principal component analysis (PCA) was performed on Perseus (MaxQuant, Munich, Germany). A 3D PCA plot was graphed using the Plotly package on R Studio. Euclidian distance was calculated between the median x, y and z coordinates for each treatment group. Linear regression analysis between Euclidean distance within the 3D PCA space and drug synergy was performed with respective combination index and ZIP synergy scores on GraphPad Prism.

#### Analysis of in vivo drug synergy

Analysis of *in vivo* drug synergy was performed using the R package invivoSyn which assesses joint action of drug combinations from PDX tumour growth curve data in mouse models which are typically limited to 4 treatment arms (Vehicle, drug A, drug B, A + B) (45, 46).

## Acknowledgments

PDX models were developed with support from Neuroblastoma Australia, or as part of the Zero Childhood Cancer Program. This study was supported by the Kids Cancer Alliance, a Cancer Institute New South Wales Translational Cancer Research Centre (JIF, TNT) and by Neuroblastoma Australia (JIF, TNT). We would like to thank the Sydney Children’s Tumour Bank Network for providing samples and related clinical information. Schematic in Figure 3B was created in BioRender. https://BioRender.com/kddoj87

## Author contributions

Conceptualization: DRC, JZRH

Methodology: JZRH, JFH, DF, AK, AFT, JIF, TNT, PT, SLL, DRC

Investigation: JZRH, MP, JFH, KHL, BN, JN, AFT, ALC, YEIO, MC, AK, SLL, DRC

Visualization: DRC, JZRH

Funding acquisition: DRC, WK, MDN, MH, JIF

Project administration: DRC

Supervision: DRC, MH, MDN, PT, TNT, JIF

Writing – original draft: DRC, JZRH

Writing – review & editing: DRC, JZRH, WK, SLL

## Data and materials availability

The full code and description of the ordinary different equation based model of the JNK network is previously published (18). All data needed to evaluate the conclusions in the paper are present in the paper and/or the Supplementary Materials.

---

**Figure S1:**
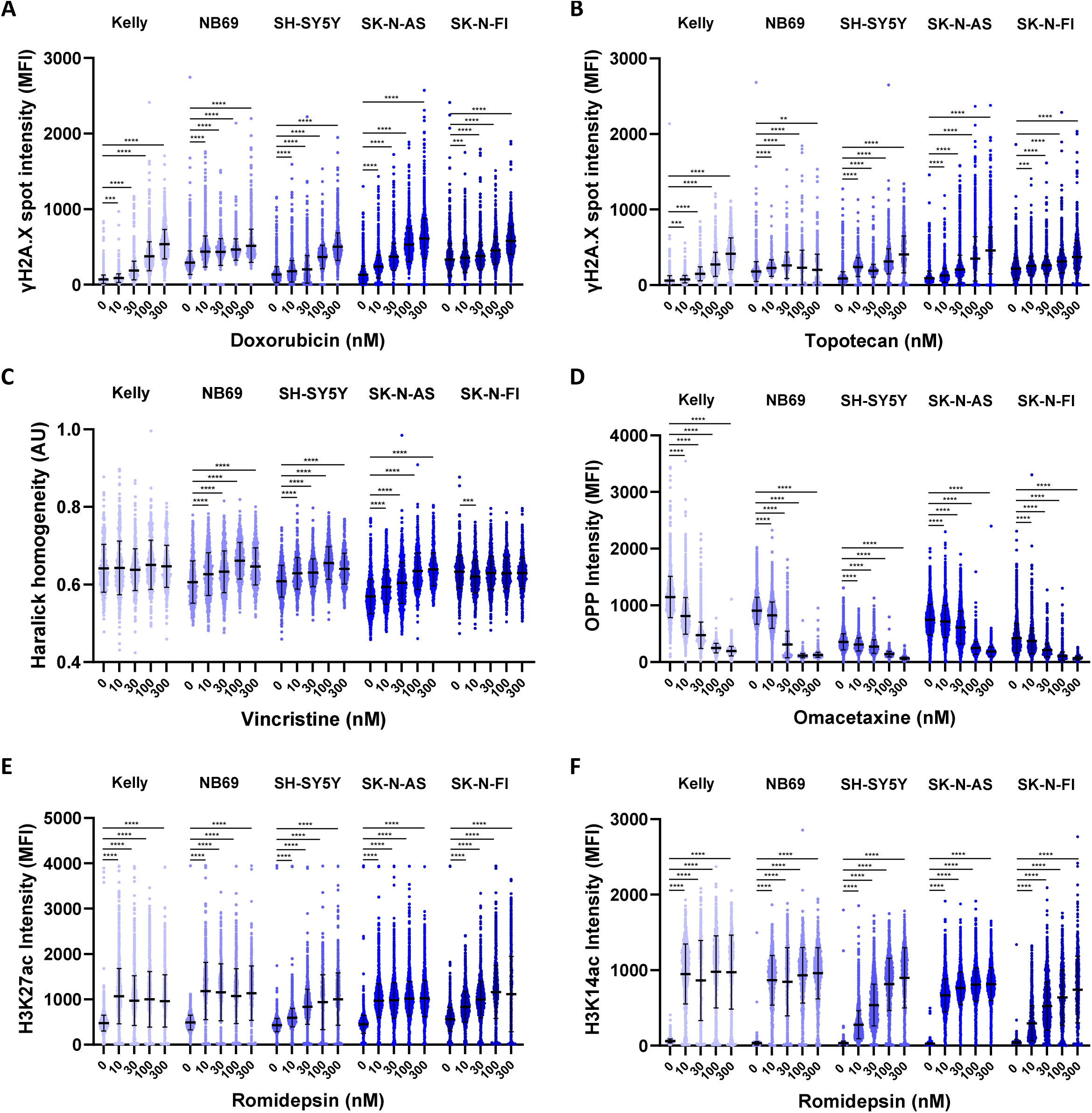
Raw data for high-content imaging of target engagement. (**A**) High-content imaging of γH2A.X antibody and DAPI staining following treatment with doxorubicin (100 nM. 24 h) or a DMSO control (Mean, n>1000). (**B**) High-content imaging of γH2A.X antibody and DAPI staining following treatment with topotecan (100 nM. 24 h) or a DMSO control (Mean, n>1000). (**C**) High-content imaging of α-tubulin antibody and DAPI staining following treatment with vincristine (300 nM. 4 h) or a DMSO control (Mean, n>1000). (**D**) High-content imaging of OPP and DAPI staining following treatment with omacetaxine (300 nM. 24 h) or a DMSO control (Mean, n>1000). (**E,F**) Quantification of relative histone acetylation at the sites indicated by high content imaging of antibody staining following romidepsin treatment at the dosages indicated (16 h. Mean, n>1000). *** p<0.001, ** p<0.01, * p <0.05.

**Figure S2:**
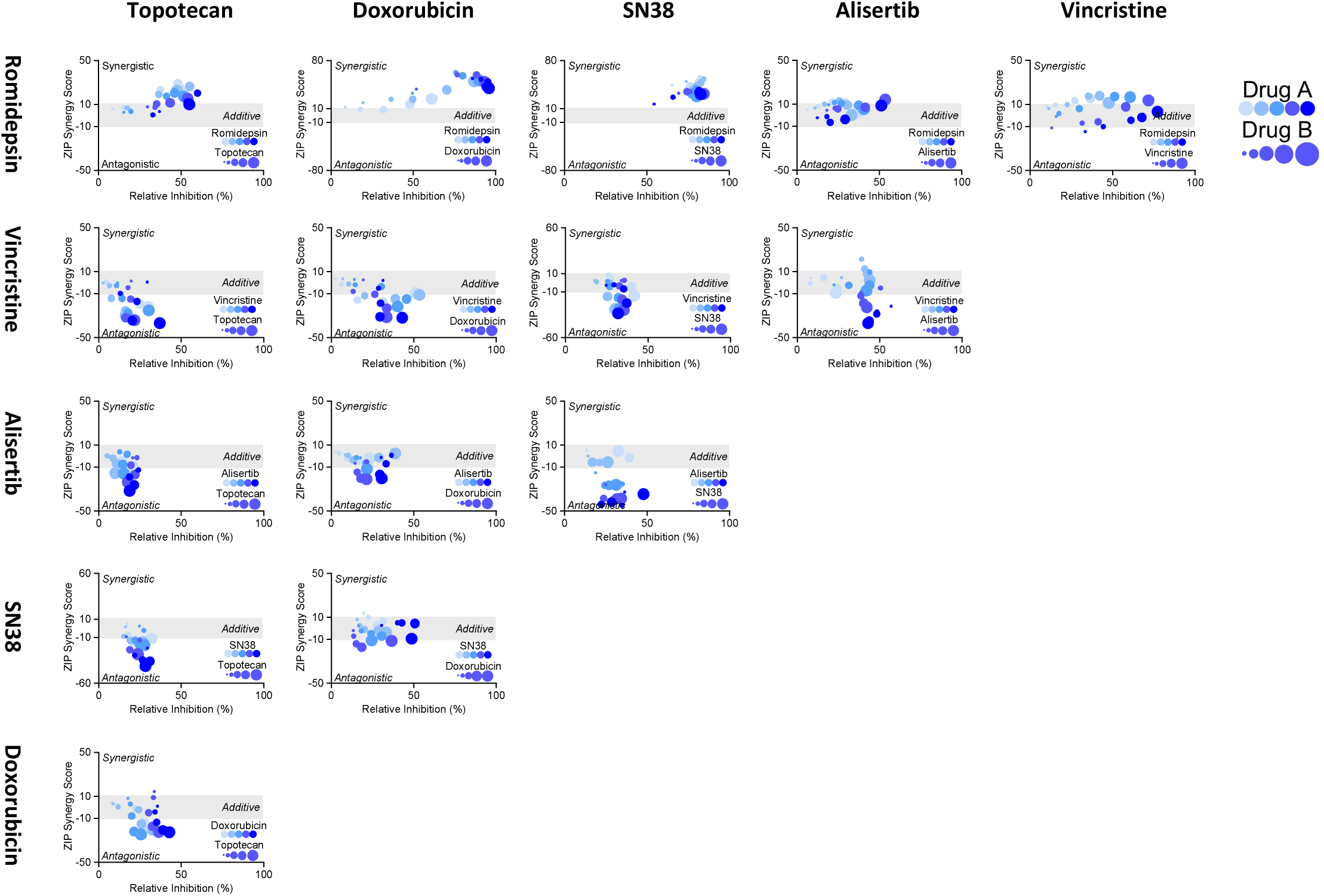
Synergy analysis of data from. Figure 5A. Quantification of ZIP synergy scores from the measurements of apoptosis inducted by each pair-wise drug combination were performed with SynergyFinder 2.0. Values lower than −10 represent and antagonistic interaction, values higher than 10 a synergistic interaction. Increasing doses of drugs on the vertical axis are indicated darker shades of blue, while increasing doses for drugs on the horizontal axis are represented by an increase in size for each data point.

**Figure S3:**
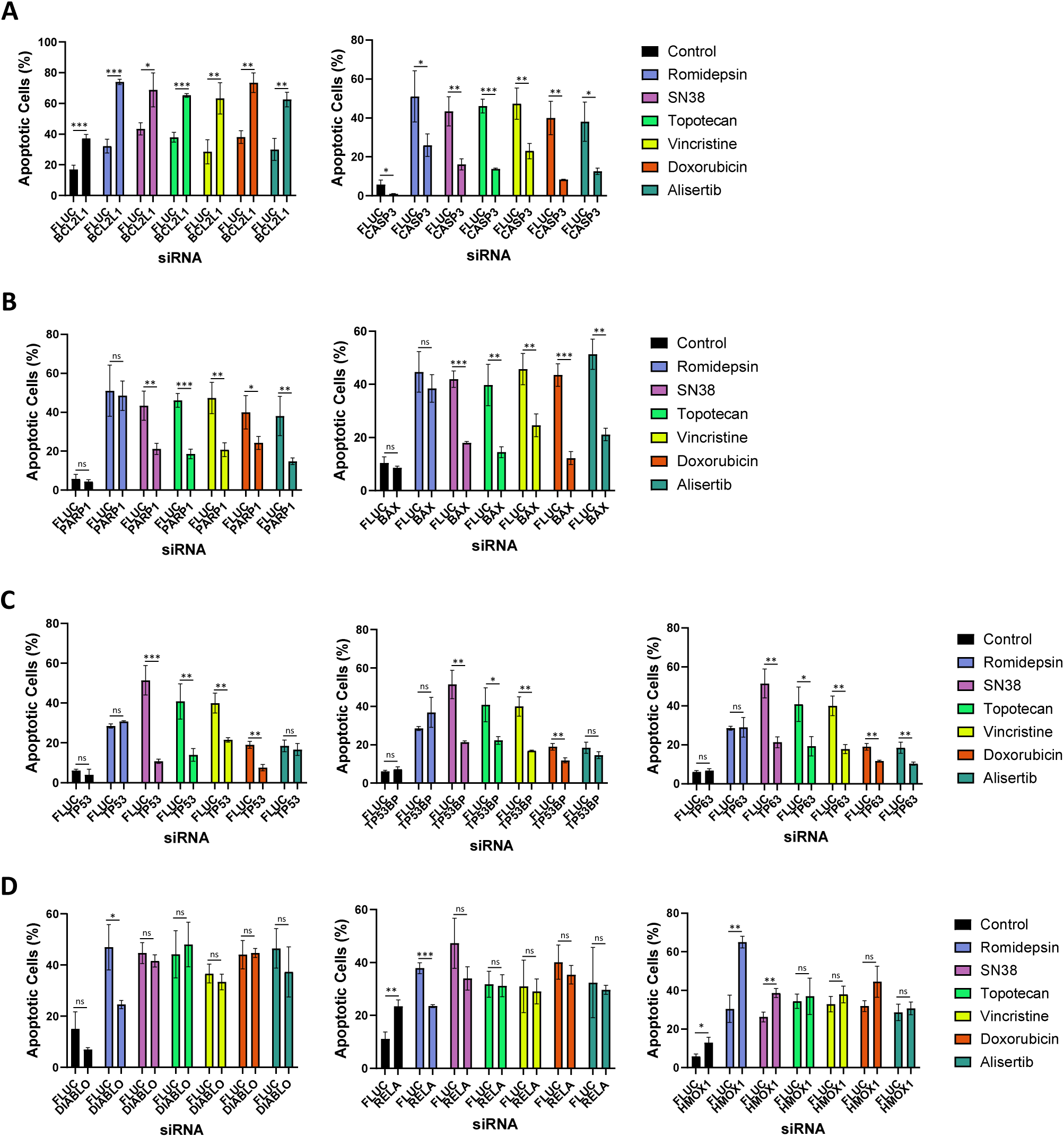
Raw data for individual genes within the apoptosis-focused functional genomics screen. (**A-D**) Raw data for the individual genes indicated from the functional genomics assay in Figure 6, combining transfection of an esiRNA library into SH-SY5Y cells followed by treatment with alisertib (50 nM), doxorubicin (50 nM), romidepsin (30nM), SN38 (15 nM), topotecan (50 nM) or vincristine (15 nM) and a high-content imaging based readout of apoptosis after staining with NucView 488 (MFI > 20). (Mean ± SD, n=3). *** p<0.001, ** p<0.01, * p <0.05.

**Figure S4:**
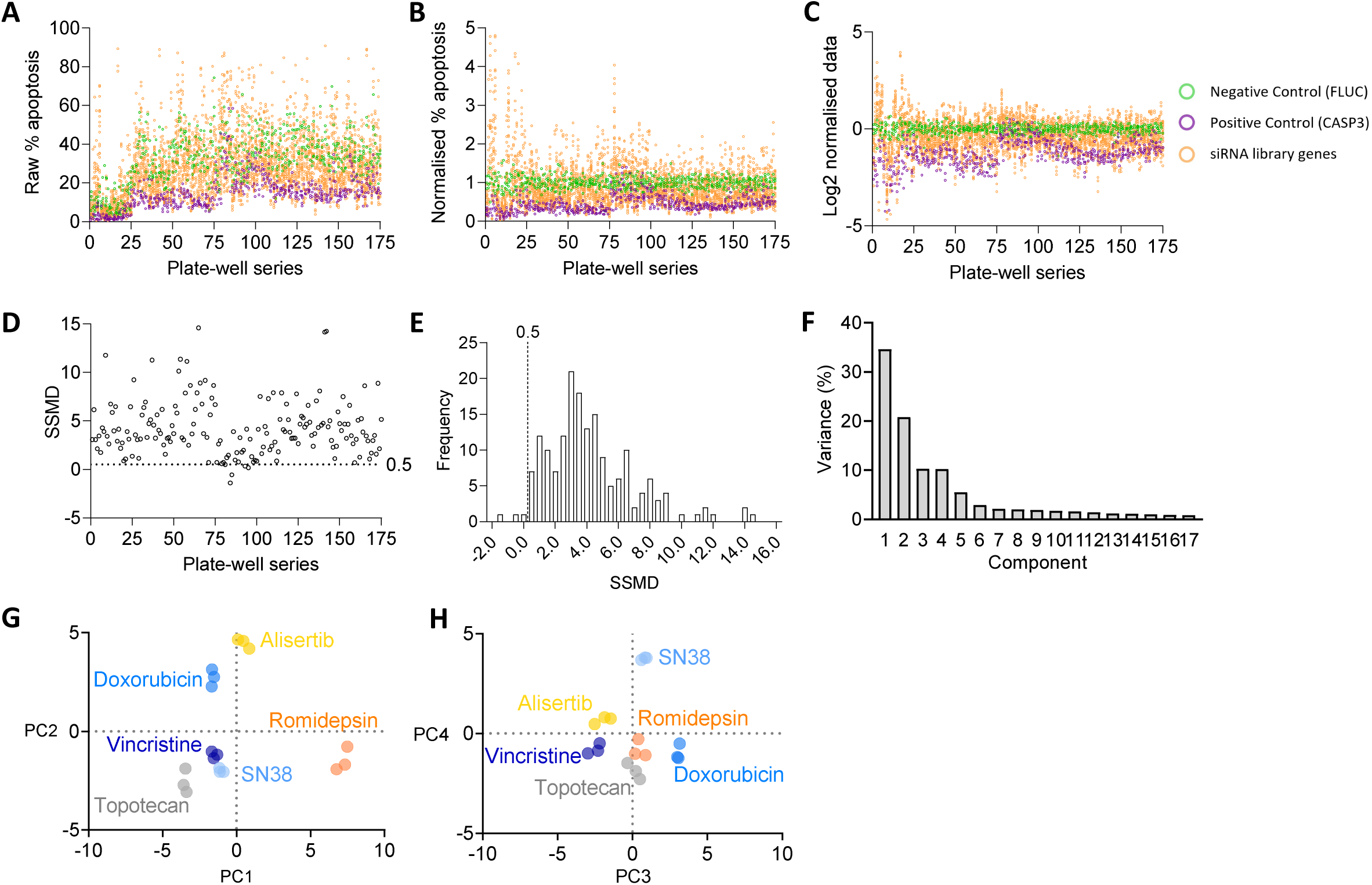
Data analysis for the functional genomics assay. (**A**) Plate-well series of the functional genomics screen data in Figure 6B. Data was collected per esiRNA, per experimental plate, in triplicate. (**B**) Plate-level data was normalised to % apoptosis of each plate’s internal negative control (FLUC). (**C**) Screen-level data was log2 transformed. (**D**) Strictly standardised mean deviation (SSMD) as a quality control metric of the functional genomics screen data, presented as a plate-well series of SSMD values generated for all 96-well plates. (**E**) A frequency distribution of SSMD scores. Each bin = 2.0 SSMD, cut-off for RNAi screen = 0.5. (**F**) Principle component analysis of the functional genomics screen dataset. Dot plots encompassing, (**G**) components 1 & 2 and (**H**) components 3 & 4.

**Figure S5:**
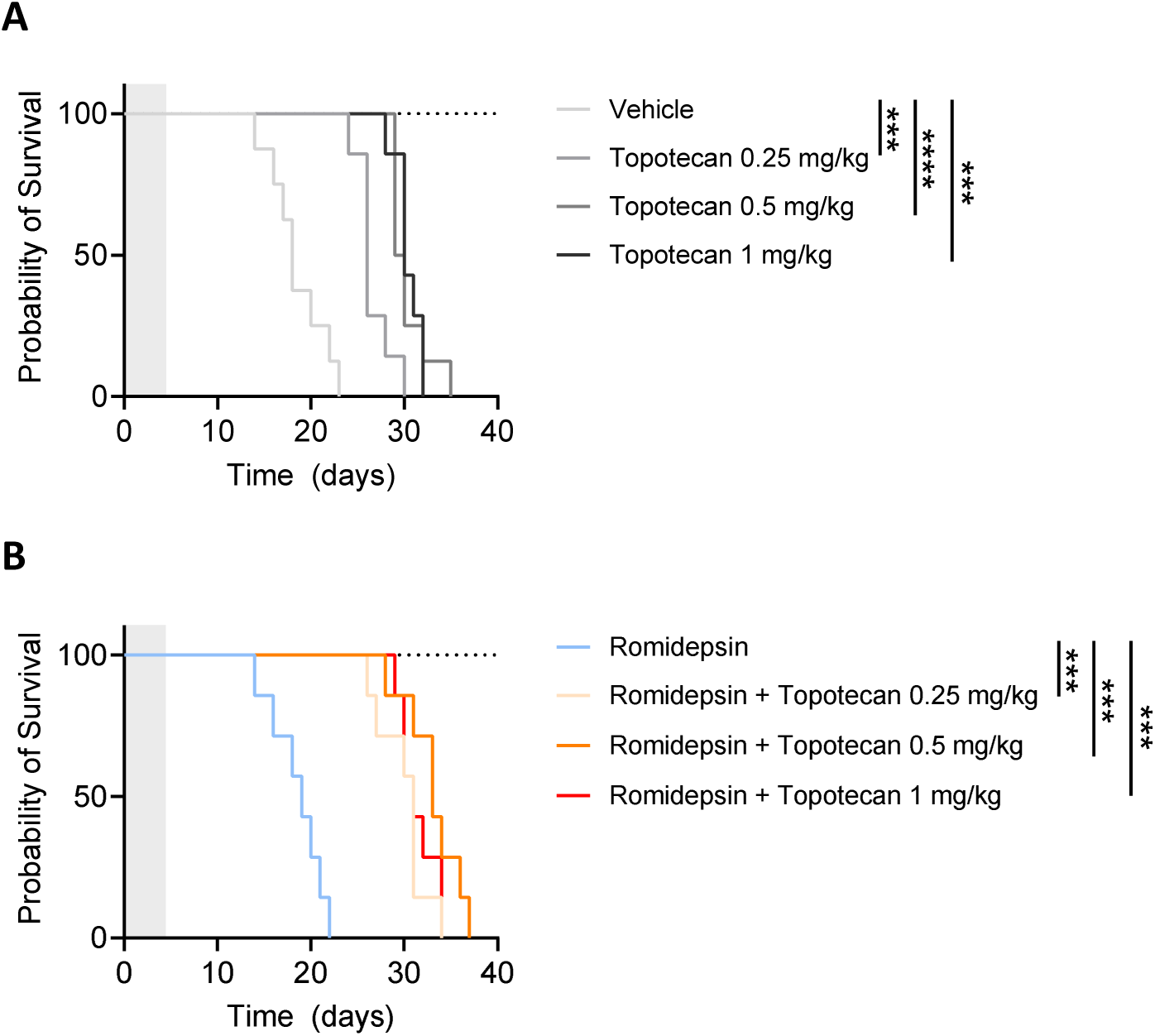
Survival analysis of topotecan and romidepsin in vivo synergy assay. NSG mice were implanted with 1×106 CCI-NB01-RPT cells, once tumours reached 150 mm3 the mice received intraperitoneal injections of the drug combinations indicated, or the relevant vehicle controls, once daily for 5 days. Tumour growth was measured every day until ethical endpoint (1000 mm3) (**A**) Survival analysis for the topotecan single-agent treatment arms. (**B**) Survival analysis for the topotecan plus romidepsin treatment arms. **** p<0,0001, *** p<0.001.

