## Supplementary Figures for "Inclusion of JNK-independent drugs within multi-agent chemotherapy improves response in relapsed high-risk neuroblastoma"

### Supplementary Figure 1

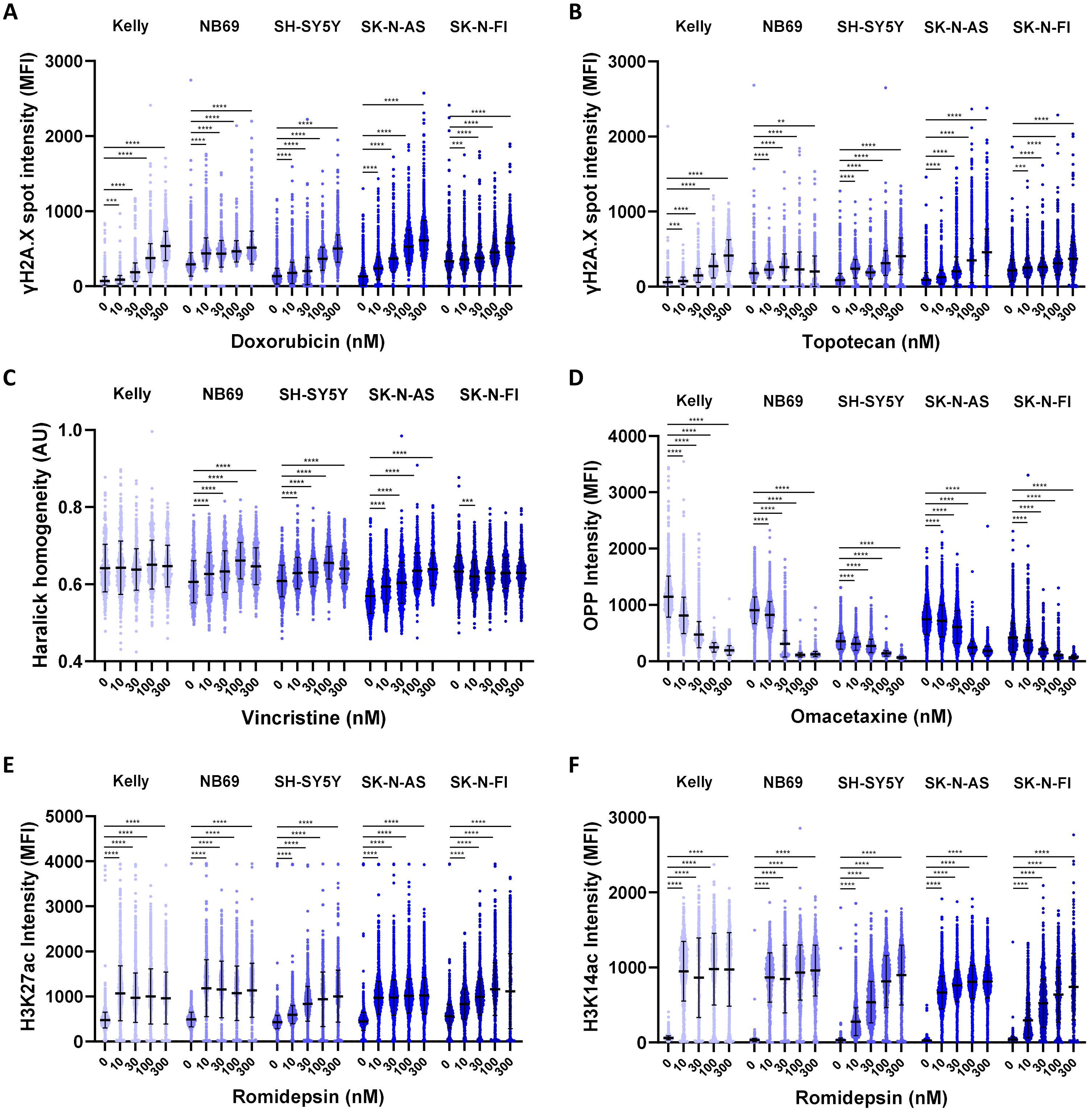


**Figure S1: Raw data for high-content imaging of target engagement.** (**A**) High-content imaging of $\gamma$H2A.X antibody and DAPI staining following treatment with doxorubicin (100 nM. 24 h) or a DMSO control (Mean, $n>1000$). (**B**) High-content imaging of $\gamma$H2A.X antibody and DAPI staining following treatment with topotecan (100 nM. 24 h) or a DMSO control (Mean, $n>1000$). (**C**) High-content imaging of $\alpha$-tubulin antibody and DAPI staining following treatment with vincristine (300 nM. 4 h) or a DMSO control (Mean, $n>1000$). (**D**) High-content imaging of OPP and DAPI staining following treatment with omacetaxine (300 nM. 24 h) or a DMSO control (Mean, $n>1000$). (**E,F**) Quantification of relative histone acetylation at the sites indicated by high content imaging of antibody staining following romidepsin treatment at the dosages indicated (16 h. Mean, $n>1000$). *** $p<0.001$, ** $p<0.01$, * $p<0.05$.

### Supplementary Figure 2

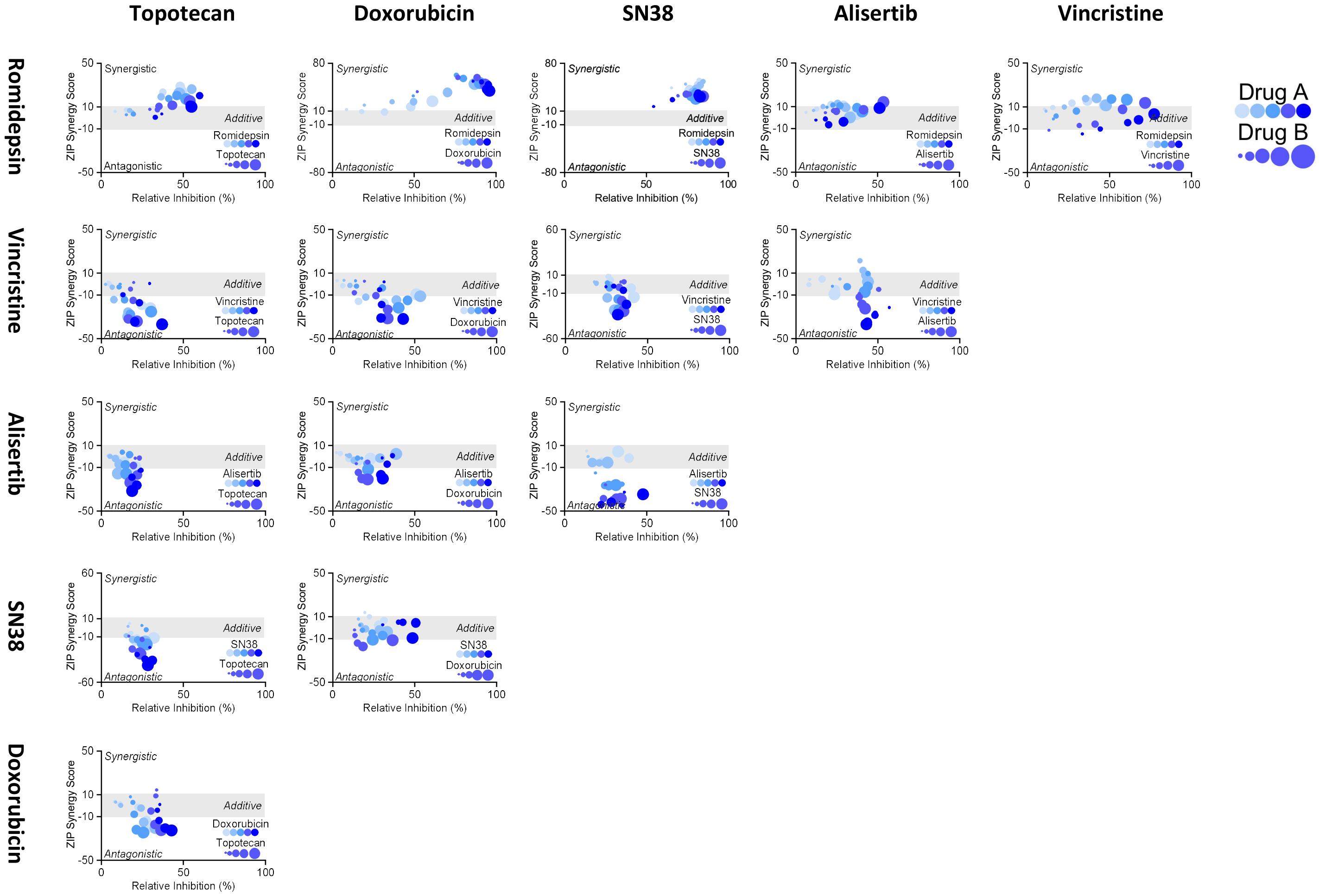


**Figure S2: Synergy analysis of data from Figure 5A.** Quantification of ZIP synergy scores from the measurements of apoptosis inducted by each pair-wise drug combination were performed with SynergyFinder 2.0. Values lower than -10 represent and antagonistic interaction, values higher than 10 a synergistic interaction. Increasing doses of drugs on the vertical axis are indicated darker shades of blue, while increasing doses for drugs on the horizontal axis are represented by an increase in size for each data point.

### Supplementary Figure 3

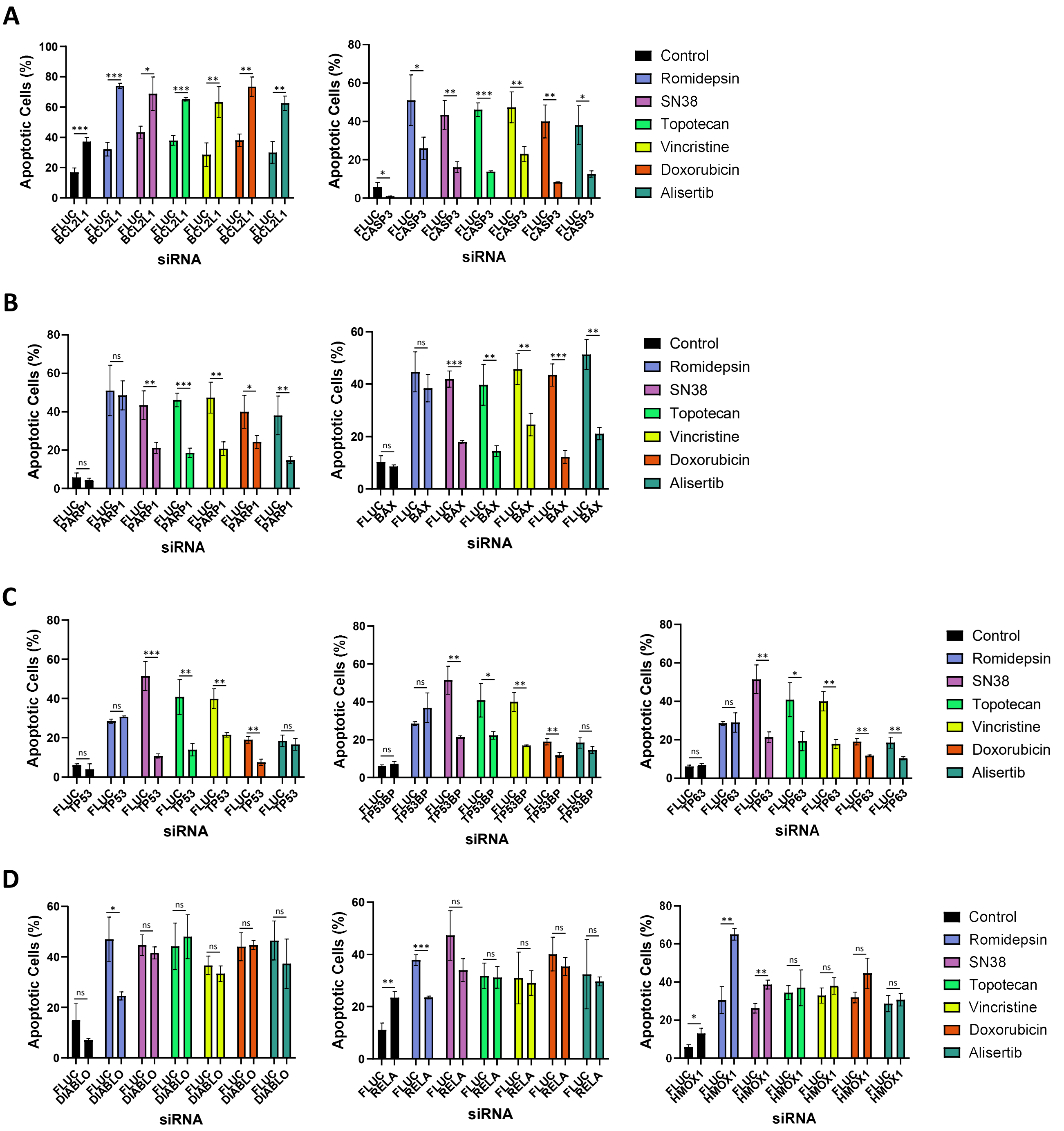


**Figure S3: Raw data for individual genes within the apoptosis-focused functional genomics screen.** (**A-D**) Raw data for the individual genes indicated from the functional genomics assay in Figure 6, combining transfection of an esiRNA library into SH-SY5Y cells followed by treatment with alisertib (50 nM), doxorubicin (50 nM), romidepsin (30nM), SN38 (15 nM), topotecan (50 nM) or vincristine (15 nM) and a high-content imaging based readout of apoptosis after staining with NucView 488 (MFI > 20). (Mean ± SD, n=3). *** $p<0.001$, ** $p<0.01$, * $p<0.05$.

### Supplementary Figure 4

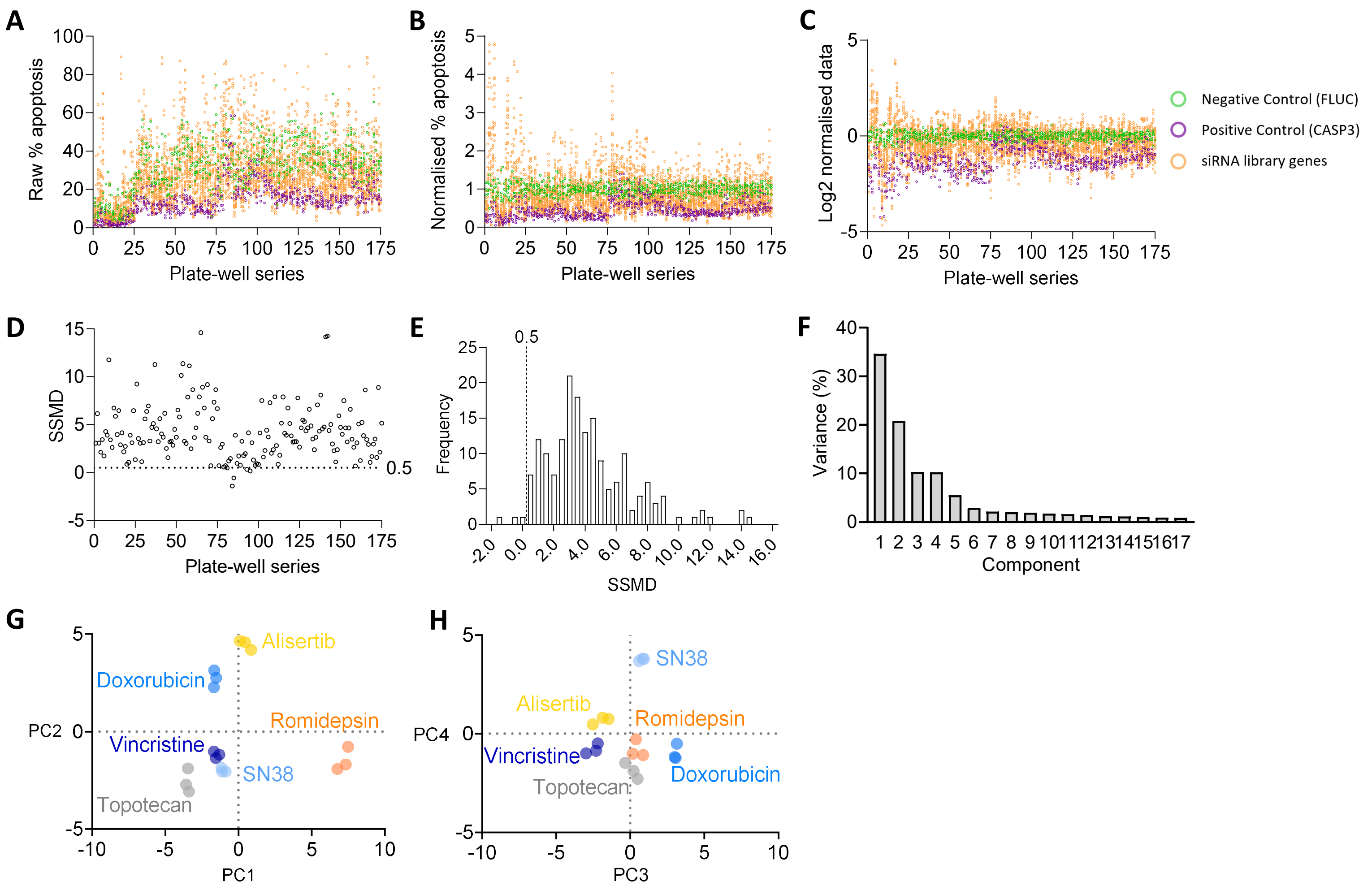


**Figure S4: Data analysis for the functional genomics assay.** (**A**) Plate-well series of the functional genomics screen data in Figure 6B. Data was collected per esiRNA, per experimental plate, in triplicate. (**B**) Plate-level data was normalised to % apoptosis of each plate's internal negative control (FLUC). (**C**) Screen-level data was log2 transformed. (**D**) Strictly standardised mean deviation (SSMD) as a quality control metric of the functional genomics screen data, presented as a plate-well series of SSMD values generated for all 96-well plates. (**E**) A frequency distribution of SSMD scores. Each bin = 2.0 SSMD, cut-off for RNAi screen = 0.5. (**F**) Principle component analysis of the functional genomics screen dataset. Dot plots encompassing, (**G**) components 1 & 2 and (**H**) components 3 & 4.

### Supplementary Figure 5

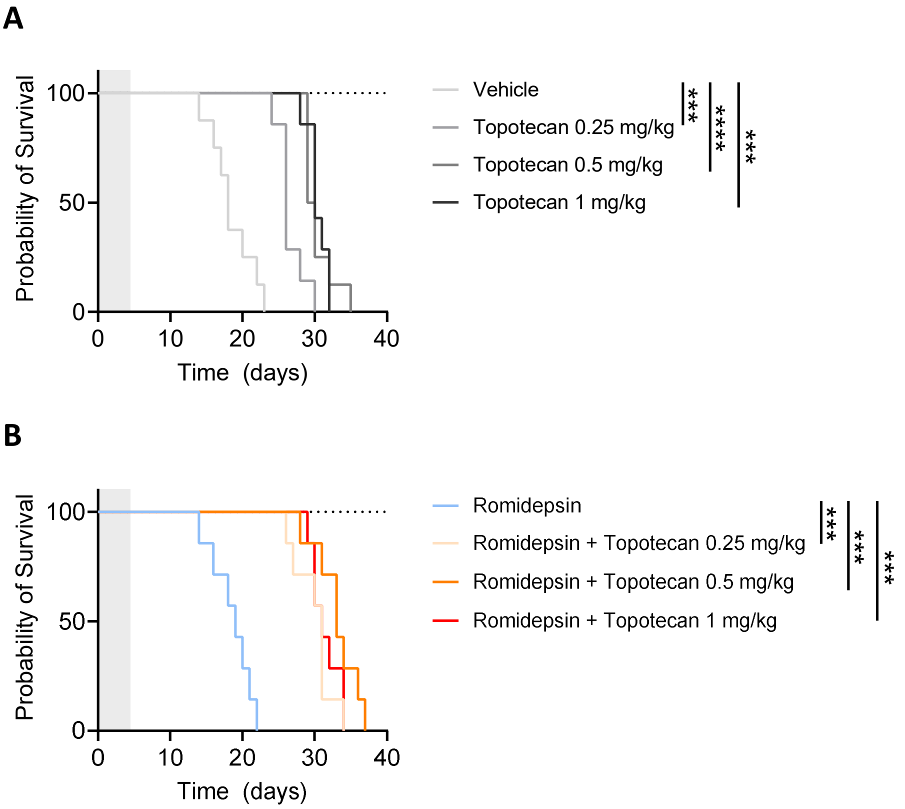


**Figure S5: Survival analysis of topotecan and romidepsin in vivo synergy assay.** NSG mice were implanted with 1x106 CCI-NB01-RPT cells, once tumours reached 150 mm3 the mice received intraperitoneal injections of the drug combinations indicated, or the relevant vehicle controls, once daily for 5 days. Tumour growth was measured every day until ethical endpoint (1000 mm3) (**A**) Survival analysis for the topotecan single-agent treatment arms. (**B**) Survival analysis for the topotecan plus romidepsin treatment arms. **** p<0,0001, *** p<0.001.
