## Supplementary Table 3 for "Inclusion of JNK-independent drugs within multi-agent chemotherapy improves response in relapsed high-risk neuroblastoma"

**Supplementary Table 3 – Reagent details**

| **REAGENT or RESOURCE** | **SOURCE** | **IDENTIFIER** |
| --- | --- | --- |
| **Antibodies** | | |
| α-JNK | Cell Signaling | Cat# 9252 |
| α-phospho-JNK T183/Y185 | Cell Signaling | Cat# 9251  (Western Blotting) |
| α-phospho-Akt S473 | Cell Signaling | Cat# 4060 |
| α-Cleaved Caspase 3 | Cell Signaling | Cat# 9661 |
| α-Cleaved Caspase 7 | Cell Signaling | Cat# 9661 |
| α-MCL-1 | Cell Signaling | Cat# 4572 |
| Acetyl-Histone H3 Antibody Sampler Kit (Lys 9, 14, 18, 27, 56 and Total Histone H3) | Cell Signaling | Cat# 9927 |
| α-MKK4 | Abcam | Cat# ab33912 |
| α-ZAK | Abcam | Cat# ab65249 |
| α-Actin (AC-15) | Merck | Cat# A1978 |
| α-MKK7 | Acris Antibodies | Cat# AM00096PU-N |
| α-Ki67 | Cell Signaling | Cat# 5315 |
| α-γH2A.X | Bioss | Cat# bs-13546R-A488 |
| **Experimental Models: Organisms/Strains** | | |
| NOD.Cg-Prkdc<scid>IL2rg<tm1Wjl>/SzJAusb | Australian BioResources |  |
| **Chemicals** | | |
| Vorinostat | Selleck Chemicals | Cat# S1047 |
| EPZ-6438 | Selleck Chemicals | Cat# S7128 |
| GSK-343 | Selleck Chemicals | Cat# S7164 |
| ABT-199 | Selleck Chemicals | Cat# S8048 |
| JNK-IN-8 | Selleck Chemicals | Cat# S4901 |
| Anisomycin | Sigma-Aldrich | Cat# A9789 |
| Vincristine | Sigma-Aldrich | Cat# V8388 |
| S63845 | MedChemExpress | Cat# HY-100741 |
| Hoechst 33342 | Thermo Fisher Scientific | Cat# 62249 |
| **Software and Algorithms** | | |
| Ordinary Different Equation based JNK Network Model | (Fey et al., 2015) |  |

| Cell line | Species | Source | Product no. | Media |
| --- | --- | --- | --- | --- |
| BE(2)-C | Human | ATCC | CRL-2268 | DMEM (Cat#11965118) 10% FCS, 1% penstrep |
| HEK-293t | Human | ATCC | CRL-3216 |  |
| Plat-E | Human | CellBioLabs | RV-101 |  |
| Kelly | Human | Riken Cell Bank | CVCL_2092 | RPMI 1640 (Cat#11875093) 10% FCS, 1% penstrep |
| NB69 | Human | Riken Cell Bank | CVCL_1448 |  |
| SH-SY5Y | Human | ATCC | CRL-2266 |  |
| NB9464 | Mouse | UCLA | - | RPMI 1640 (Cat#11875093) 10% FCS, 1% non-essential amino acids, 1% sodium pyruvate, 1% penstrep |
| SK-N-AS | Human | ATCC | CRL-2137 | DMEM (Cat#11965118) 10% FCS, 1% non-essential amino acids, sodium pyruvate, 1% penstrep |
| SK-N-FI | Human | ATCC | CRL-2142 |  |
| IMR32 |  |  |  |  |
